# IL-21 and IFN-alpha have both opposite and redundant role on human innate precursors and memory B-cell differentiation

**DOI:** 10.1101/2021.03.31.437810

**Authors:** Marina Boudigou, Magalie Michée-Cospolite, Patrice Hémon, Alexis Grasseau, Christelle Le Dantec, Emmanuelle Porchet, Christophe Jamin, Valérie Devauchelle, Olivier Mignen, Divi Cornec, Jacques-Olivier Pers, Laëtitia Le Pottier, Sophie Hillion

## Abstract

Immunological memory is essential for effective immune protection upon antigen rechallenge. Memory B cells encompass multiple subsets, heterogeneous in terms of phenotypes, origins and precursors, anatomical localization, and functional responses. B-cell responses are conditioned by micro-environmental signals, including cytokines. Here, we analyzed *in vitro* the effects of two cytokines implicated in B-cell differentiation, interferon-alpha (IFN-α) and interleukin (IL)-21, on the early functional response of four different mature B-cell subsets (IgD^-^ CD27^-^ naive, IgD^+^ CD27^+^ unswitched, IgD^-^ CD27^+^ switched and double-negative B cells). The dual response of naive and memory B cells to IL-21 allowed us to uncover a unique IgD^+^ CD27^-^ CD10^-^ B-cell population (referred to as NARB^+^) characterized by the expression of marginal zone B-cell markers CD45RB and CD1c. Similar to memory B cells, NARB^+^ cells were in a pre-activated state, allowing them to rapidly differentiate into plasmablasts upon innate signals while maintaining their susceptibility to IL-21 activation-induced apoptosis as observed for the naive compartment. Both in-depth phenotypic analysis of circulating B cells, and identification of these cells in spleen, tonsil and gut-associated lymphoid tissues, supported that NARB^+^ are uncommitted precursors of human marginal zone B cells.

## INTRODUCTION

Memory B cells (MBC) and antibody (Ab) secreting cells (ASC) are the key players of humoral immunity resulting in the clearance of pathogens and the establishment of durable immune protection and supporting successful vaccination strategies (1). However, the persistence of autoreactive memory B-cell clones could sustain chronic and autoreactive inflammation leading to systemic autoimmune diseases (2). Some inherent and stable memory functions include long lifespan, high sensitivity to low doses of antigen, quick and robust proliferation, and rapid differentiation into plasma cells highly contrast with those from naive inexperienced B cells (3, 4). Accumulated data have demonstrated the high degree of heterogeneity of subsets and their progenitors with ambivalent equivalence in humans (5). Activated B cells that make cognate interactions in secondary lymphoid organs differentiate along either a follicular or extra-follicular (EF) pathways. In the follicular pathway, antigen-activated B cells interact with T cells and form the germinal center (GC), where they undergo somatic hypermutation, selection, and eventually differentiate toward high affinity long-lived plasma cells or IgG^+^ memory B cells. The EF differentiation of B cells is described as very heterogeneous depending on the tissue localization and subsequently on the nature of the availability of T-cell help as described in the lymph node (6). In the gut or the spleen, this pathway is mostly T-cell independent but relies on signals provided by innate cells (7). However, the strength of antigen (Ag) recognition appears as a common crucial regulator (8). The prototype of this pathway is represented by marginal zone B-cell (MZB) differentiation, leading to the production of short-lived plasmablasts (PBs) involved in bacterial protection and the immune regulatory response (9).

The cytokines IL-21 and IFN-α are major actors involved in the control of B-cell responses. In secondary organs, the production of IL-21 by follicular helper T cells (Tfh) not only initiates affinity maturation in the GCs and isotype switching but also promotes cell proliferation and plasma cell differentiation (10–12). IFN proteins are highly produced by plasmacytoid dendritic cells upon viral aggression and are involved in the development of protective immunity mostly against viruses (13–15). Interestingly, IL-21 and IFN-α signaling have shared involvement in the pathogenesis of chronic and autoimmune diseases as well as in uncontrolled exacerbated inflammation (16–18). However, the role of IL-21 and IFN-α role in influencing the reactivation of MBCs or initiating the innate B-cell differentiation pathway is not well understood and often analyzed separately, especially in humans (19, 20). In the present report, we were interested in exploring the role of both cytokines in the initial events that cause activation and plasma cell differentiation from different human mature B-cell progenitors. Thus, we thus used a canonical gating strategy with IgD and CD27 to sort the four main mature B-cell populations. We then analyzed the early response of B cells to IL-21 and IFN-α signaling following BCR and Toll-like receptor (TLR) stimulation. We showed that both signaling pathways induced different B-cell fates, depending on their maturation status. Furthermore, we uncovered a mature B-cell population IgD^+^ CD27^-^ CD45RB^+^ CD10^-^ (referred to as NARB^+^) harboring highly versatile “multilineage” differentiation potential either able to differentiate into IgM producing PB upon innate signals or adopt pre-GC features in the presence of IL-21.

Phenotypically, NARB^+^ cells share some attributes with IgD^+^ CD27^+^ MZB cells but retain a unique naive migratory pattern and are sensitive to IL-21 activation-induced apoptosis. This population differed from Ag-inexperienced naive B cells by exhibiting a pre-activated phospho-signaling response to BCR/TLR stimulation. The co-stimulation with IFN-α but not with IL-6 or IL-10 overtook the IL-21 dependent inhibition of the differentiation pathway. This functional plasticity towards its cytokine environment contrasted with the stable and redundant responses observed in committed MBC subsets. Therefore, this subset could represent an important crossroad in human between the follicular and marginal zone developmental pathways.

## MATERIALS AND METHODS

### Sample collections

Blood samples from healthy donors were collected. Routine plateletpheresis was performed with the Trima Accel device at the French Blood Establishment (EFS). After donation, tubing of the leukoreduction system chamber (LRSC) was sealed before removing the collection set from the Trima Accel system. The LRSC were kept at 4°C until cell recovery (within 18 hours).

### Cell isolation

To isolate Peripheral Blood Mononuclear Cells (PBMCs) from Trima Accel devices, residual human blood from the LRSC was harvested and the LRSC was rinsed with phosphate-buffered saline (PBS). PBMCs were isolated by density gradient centrifugation on Lymphocyte Separating medium, Pancoll human (PAN Biotech). CD19^+^ B cells were purified from human PBMCs using the REAlease® CD19 Microbead Kit (Miltenyi Biotec) according to the manufacturer’s recommendations with purity greater than 98 %.

### Cell sorting

After isolation, B cells were stained for 30 min at 4°C into the dark with fluorescein isothiocyanate (FITC)-conjugated anti-IgD (IA6-2), phycoerythrin (PE)-conjugated anti-CD10 (ALB1), phycoerythrin linked to cyanin 5.5 (PC5.5)-conjugated anti-CD27 (1A4CD27), allophycocyanin (APC) linked to Alexa Fluor 700 (AAF700)-conjugated CD19 (J3-119) antibodies. CD10-positive B cells, which correspond to transitional subset, were excluded from the cell sorting. Naive (NA, IgD^+^ CD27^-^ CD10^-^), unswitched memory (USM, IgD^+^ CD27^+^ CD10^-^), switched memory (SM, IgD^-^ CD27^+^ CD10^-^) and double-negative B cells (DN, IgD^-^ CD27^-^ CD10^-^) were sorted. For sorting of CD45RB^+^ and CD45RB^-^ NA B cells, FITC-conjugated anti-CD45RB (MEM-55, Immunotools), combined with PE-conjugated anti-CD10, PC5.5-conjugated anti-CD27, APC-conjugated anti-IgD (IA6-2) and AAF700-conjugated anti-CD19 antibodies, was used. Cell sorting was performed using a MoFlow XDP cell sorter (Beckman Coulter). The purity of B-cell subsets was typically greater than 97 %.

### Ex vivo B-cell phenotyping by Flow Cytometry

For B-cell phenotyping, the following antibodies were used (Supplementary Table 1). For ABCB1 transporter activity analysis, cells were then washed and incubated at 37°C for 30 min with 200 nmol/L MitoTracker Green FM probes (Molecular Probes, Eugene, Ore). Samples were acquired with a Navios cytometer (Beckman Coulter).

### Cell culture

Sorted cells were cultured in 96-well plates (Falcon) at 2×10^5^ cells per 200 µL/wells in complete medium (RPMI 1640 medium (Sigma-Aldrich) supplemented with 10 % of heat-inactivated Fetal Calf Serum (FCS) (BD, Biosciences), 4 mM L-Glutamine (Gibco) and penicillin (200 U/mL) and streptomycin (100 µg/mL)). B cells were stimulated for 84 h (3.5 days), in the presence of the AffiniPure Goat anti-Human IgG+IgM (H+L) (2 µg/mL), the AffiniPure F(ab’)2 Fragment Goat Anti-Human Serum IgA, α Chain Specific (2 µg/mL; Jackson ImmunoReasearch Laboratories), CpG oligodeoxynucleotide (ODN-2006) (0,25 µM; InvivoGen), recombinant human Interleukin-2 (rh IL-2) (20 ng/mL, ImmunoTools) subsequently referred to as primed stimulation. Recombinant human Interferon-alpha 2a (rh IFN-α2a) (10 µg/mL; ImmunoTools), or recombinant human IL-21 (40 ng/mL; Immunotools) were added to the stimulation. For kinetics studies, B cells were stimulated with or without IL-21 for 12 h (0.5 days), 1.5 days, 2.5 days and 3.5 days.

### Apoptosis and differentiation rescue assays

Viability of stimulated B cells were assessed by flow cytometry using FITC-labeled annexin V and propidium iodide (PI) (Biolegend). For apoptosis assays by live-imaging, IncuCyte^®^ Annexin V Green Reagent was added directly in the medium at the dilution 1:200 and B cells were imaged every hour for 3.5 days using IncuCyte^®^ Live-Cell Analysis System (Sartorius).

For rescue assays, sorted B cells were cultured with rh IL-21 as previously described and with Human CD40-Ligand Multimer Kit (1 µg/mL, Miltenyi Biotec), rh IL-6 (20, 100 or 1000 ng/mL, Immunotools), rh IL-10 (10, 100 or 1000 ng/mL, Immunotools), rh BAFF (50 ng/mL, Immunotools), rh IFN-α2a (10 ng/mL) or with the Pan Caspase Inhibitor Z-VAD-FMK (100 µM, BD Biosciences).

### Analysis of the differentiation of cultured B cells by Flow Cytometry

After culture, cultured B cells were harvested, pelleted and the supernatant was collected. Cells were then stained with PE anti-CD11c, PC5.5 anti-CD27, APC-AF700 anti-CD19, APC-AF750 anti-CD38 (Beckman Coulter).Intracellular staining for transcription factors was carried out after cell fixation and permeabilization using the Transcription Factor Buffer Set (BD Biosciences) according to the manufacturer’s instructions. Transcription factor expression was analyzed using PE anti-IRF8 (REA516), PE Vio770 anti-IRF4 (REA201), APC anti-BLIMP1/PRDM1 (646702), BV421 anti-T-BET (4B10, Biolegend). Stained B cells were analyzed using a Navios cytometer (Beckman coulter). The phenotype of the cells was analyzed using Kaluza Flow Cytometry Analysis and FlowJo software.

### Phosphoprotein measurement

Purified total B cells were first stained with FITC-conjugated anti-CD45RB, PC5.5-conjugated anti-CD27, APC-conjugated anti-IgD, AAF700-conjugated anti-CD19 and AAF750-conjugated anti-CD38 antibodies for 30 min at 4°C into dark. After surface staining, cells were stimulated following the primed stimulation with or without rh IL-21 or rh IFN-α2a (as described above) during 5 to 30 min. Unstimulated B cells were used as control. Cells were then fixed in 1.5 % Paraformaldehyde 10 min at room temperature and permeabilized with 100 % ice-cold methanol for 30 min at 4°C. The expression of phosphorylated proteins was measured by flow cytometry using a Navios cytometer with VioBlue (VB)-conjugated anti-pSTAT1 (pY701, REA159), Brilliant Violet 421 (BV421)-conjugated anti-pSTAT3 (pY705, 13A3-1) or BV421-conjugated anti-pAKT (pS473, M89-61) (Biolegend) Abs.

### Proliferation assays

Sorted B-cell subsets were labeled with CellTrace Violet reagent (Thermo Fischer Scientific) for 10 min at 37°C prior to the primed stimulation as described above. B-cell proliferation was evaluated by using Navios cytometer and analyzed with FlowJo’s Proliferation software.

### Antibody production measurement by FluoroSpot

After 3.5 days of stimulation, the levels of total IgM, IgG, and IgA production by each sorted B-cell subset were evaluated by using the Human IgG/IgM and IgG/IgA FluoroSpot kits (MABTECH), according to the manufacturer’s instructions. 15,000 pre-stimulated B cells were incubated by wells during 20 h before revelation. Spots were counted with an Elispot Reader system (Autoimmun Diagnostika GmbH).

### Intracellular cytokine detection

For analysis of the intracellular cytokine production, 50 ng/mL Phorbol 12-Myristate 13-Acetate (PMA, Sigma Aldrich), 1 µg/mL ionomycin (Sigma Aldrich) and 10 µg/mL Brefeldin A (Selleckchem) were added for the last 5 hours of culture. Cells were then fixed and permeabilized using Cyto-Fast Fix/perm Buffer Set (Biolegend) according to the manufacturer’s instructions and stained with PE anti-IL-6 and PE-Cy7 anti-IL-10 (JES3-9D7) Abs (Biolegend).

### Blood sample staining and CYTOF acquisition

Whole blood from healthy controls was analyzed using the Maxpar® Direct™ Immune Profiling Assay™ (Fluidigm) according to manufacturer’s instructions on a CYTOF Helios system (Fluidigm). The following antibodies were added to the initial cocktail: CD24-169Tm (ML5, Fluidigm), CD21-162Dy (BL-13, Miltenyi Biotec), CD11b-209Bi (ICRF44, Fluidigm), IgM-175Lu (PJ2-22H3, Miltenyi Biotec), T-BET-165Ho (4B10, Biolegend), FoxP3-159Tb (259D/C7, Fluidigm) and CD45RB-142Nd (MEM-55, Immunotools). Briefly, 270 µl of blood were incubated in dry antibody tube with the additional antibodies (CD24, CD21, CD11b, IgM and CD45RB). After 30 min erythrocyte were lysed by addition of 250 µl of Cal-lyse and washed. Then cells were fixed and permeabilized using the Maxpar® Nuclear Antigen Staining Buffer Set Cells before incubation with nuclear antibodies (T-BET and Foxp3). Then cells were fixed in 1.6% paraformaldehyde and incubated with 125mM Cell-ID™ Intercalator-Ir. Sample acquisition was performed on a Helios system utilizing CyTOF® Software using the Maxpar Direct Immune Profiling Assay template with additional channels. Samples were acquired in a cell acquisition solution (CAS) containing 0.1× EQ™ beads (Fluidigm) to normalize data.

### Tissue staining and imaging mass cytometry image acquisition

Formalin-fixed paraffin-embedded tissue sections of 4[box]μm thickness from gut, tonsil and spleen, were cut onto glass slides. Sections were de-paraffinized with xylene and carried through sequential rehydration from 100% Ethanol to 70% Ethanol before being transferred to Tris-buffered saline (TBS). Heat-induced antigen retrieval was performed in a water bath at 95°C for 30 min in Tris/EDTA buffer (10mM Tris, 1mM EDTA, pH9). Slides were cooled to room temperature (RT) and were subsequently blocked with PBS+3%BSA for 30 min at RT. Each slide was incubated with 100 μ the antibody cocktail (Supplementary Table 2) overnight at 4°C. Then, slides were washed 3 times with PBS and labeled with 1:500 dilution of Intercalator-Ir (Fluidigm) in TBS for 2 min at RT. Slides were briefly washed with H2O and air dried before imaging mass cytometry (IMC) acquisition. The IMC was purchased from Fluidigm (Fluidigm, Hyperion Imaging System™). Data were acquired on a Hyperion imaging system coupled to a Helios Mass Cytometer (Fludigm), at a laser frequency of 200[box]Hz and laser power of 3[box]dB. For each recorded region of interest (ROI), stacks of 16-bit single-channel TIFF files were exported from MCD binary files using MCD™ Viewer 1.0 (Fluidigm). Cell-based morphological segmentation was carried out by using supervised pixel classification by Ilastik (64) to identify nuclei, membrane and background, and used CellProfiler (65) to segment the resulting probability maps. Inputs of 16-bit TIFF images with their corresponding segmentation mask were uploaded in histoCAT (66) to open a session data analysis. Dimensionality reduction and unsupervised FlowSOM clustering for 16-bit single images were performed using Cytobank on FCS.files.

### RNA sequencing

Total RNA from *ex vivo* sorted B cells (NA, SM and DN) and after culture (Primed, IFN-α and IL-21) was extracted using RNA isolation kits (Norgen Biotek) according to the manufacturer’s instructions. Libraries were generated by using the Qiaseq RNA library kit (Qiagen) according to manufacturer’s instructions. The sequencing analysis was realized by Macrogen society. Fastq reads were aligned using Partek® Flow® Genomic analysis software with STAR aligner and hg38-GENCODE Genes (release 31) as transcript model. Differential expression analysis was performed using the EdgeR package. We treated the PCA loadings onto each of the genes as expression data to run pathway analysis with the PGSEA package. For each pathway, the PAGE algorithm performed one-sample t-test on each gene set in the biological processes branch of Gene Ontology (GO). The adjusted P-values were then used to rank the pathways for each of the first five principal components. The following R code performs PCA and PGSEA analyses: pca.object <- prcomp(t(x)); pg = PGSEA (pca,cl=GeneSets(),range=c(15, 2000), p.value=TRUE, weighted=FALSE). We used the top 500 genes in hierarchical clustering using the heatmap.2 function. The data was centered by subtracting the average expression level for each gene.

### Real-time PCR

From 120 to 500 ng of total RNA were reverse-transcripted with superscript IV reverse transcriptase (Thermo Fisher Scientific) according to the manufacturer’s instructions. The gene expression was analyzed with the TaqMan™ Gene Expression Master Mix and the following probes: *GAPDH* (Hs02758991), *BCL2* (Hs04986394), *IL6* (Hs00985639), *IL6R* (Hs01075666), *IL10* (Hs00174086) and *TNFSF13B* (Hs00198106). Real-time PCR were performed using The QuantStudio6™ real-time PCR device (QuantStudio™ real-time PCR software). The mRNA relative abundances in real-time PCR were obtained by using the MCMC.qPCR package.

### Statistics and Data visualization

For *in vitro* experiments, statistical differences were evaluated using the paired two-tailed Wilcoxon signed-rank t-test or repeated measures (RM) one-way ANOVA using Tukey’s correction for multiple comparisons when analyzing more than three groups. All statistical analyses were performed using GraphPad Prism version 8.4.3 and 9 software.

## RESULTS

### Activation, rapid differentiation and immunoglobulin production upon stimulation define functional B-cell subsets

To investigate the initial activation and differentiation response of the different circulating mature B-cell subsets, we sorted the four canonical mature B-cell populations in CD10-negative cells according to IgD and CD27 expression. The expression of both markers was widely used to phenotype and follow the mature human B-cell population in the literature (21, 22) and delineates naive B cells (NA, IgD^+^ CD27^-^), unswitched MBCs (USM, IgD^+^ CD27^+^), switched MBCs (SM, IgD^-^ CD27^+^), and double-negative cells (DN, IgD^-^ CD27^-^) (Supplementary Figure 1A). We analyzed activation and differentiation capacities of the different subsets after activation (anti-BCR, CpG, and IL-2) (23) during a short incubation period (three days) that was inferior to previous culture conditions required for the *in vitro* class switch recombination (24). In subsequent experiments, we referred to this stimulation as priming activation. We first followed B-cell activation and differentiation by assessing the expression of CD38 and CD27 in culture to classify resting B cells (CD38^-^ CD27^-^), activated B cells (Act, CD38^+^ CD27^-^), intermediate (Int, CD38^+/-^ CD27^+^) and plasmablasts (PBs, CD38^high^ CD27^high^) (Figure 1A) adapted from previous studies (18, 25, 26). As expected, PBs and Int cells were significantly enriched in cultures derived from CD27^+^ MBCs compared to those from NA and DN cells, which in contrast presented an activated phenotype with enrichment of CD38^+^ CD27^-^ cells. However, we could detect that 25.9% (± 4.8) of CD38^+^ CD27^+^ cells originated from the DN population and more surprisingly 14.2% (± 1.8) from the NA population. To extend these phenotypic observations, we assessed the IgM, IgA, and IgG production by dual-color ELISPOT in activated sorted B-cell subsets (Figure 1B). Detection of isotype-specific PB confirmed that the culture model does not allow for the class-switching of IgD-positive and IgM-positive cells from NA and USM cells, supporting the observation of the rapid differentiation of MBC progenitors. Although the SM population presented an increase in PB and CD27^+^ cells compared with cells originating from the DN population, the percentage of ASC (all isotypes) was very close in both populations (13.6 *vs* 10.4). Moreover, there was a significant amount of IgM-secreting cells originating from the NA population which is usually positioned as a non-secreting subset. Therefore, to understand these potent discrepancies between the observed differentiated phenotype and the functional ability of the different subsets to secrete Ab, we assessed the intracellular expression of three key molecular drivers of B-cell differentiation BLIMP1 (encoded by the *PRDM1* gene), IRF4 and IRF8 (27) (Figure 1C). We robustly detected the induction of IRF4^high^ IFR8^low^ cells expressing the PB marker BLIMP1 with no significant difference between the SM, DN, and USM populations regardless of CD27 expression. In agreement with the ELISPOT assays. NA stimulated cells generated 5.5% (± 1.4) of BLIMP1^+^ IRF4^+^ IRF8^low^ cells that did not acquire a typical CD27^+/high^ pre-PB/PB phenotype. Moreover, the detection of differentiated cells with functional Ig production derived from NA cells might indicate the existence of a precursor IgD^+^ CD27^-^ B-cell subset with the ability to rapidly differentiate into IgM^+^ PB.

**Fig. 1.**
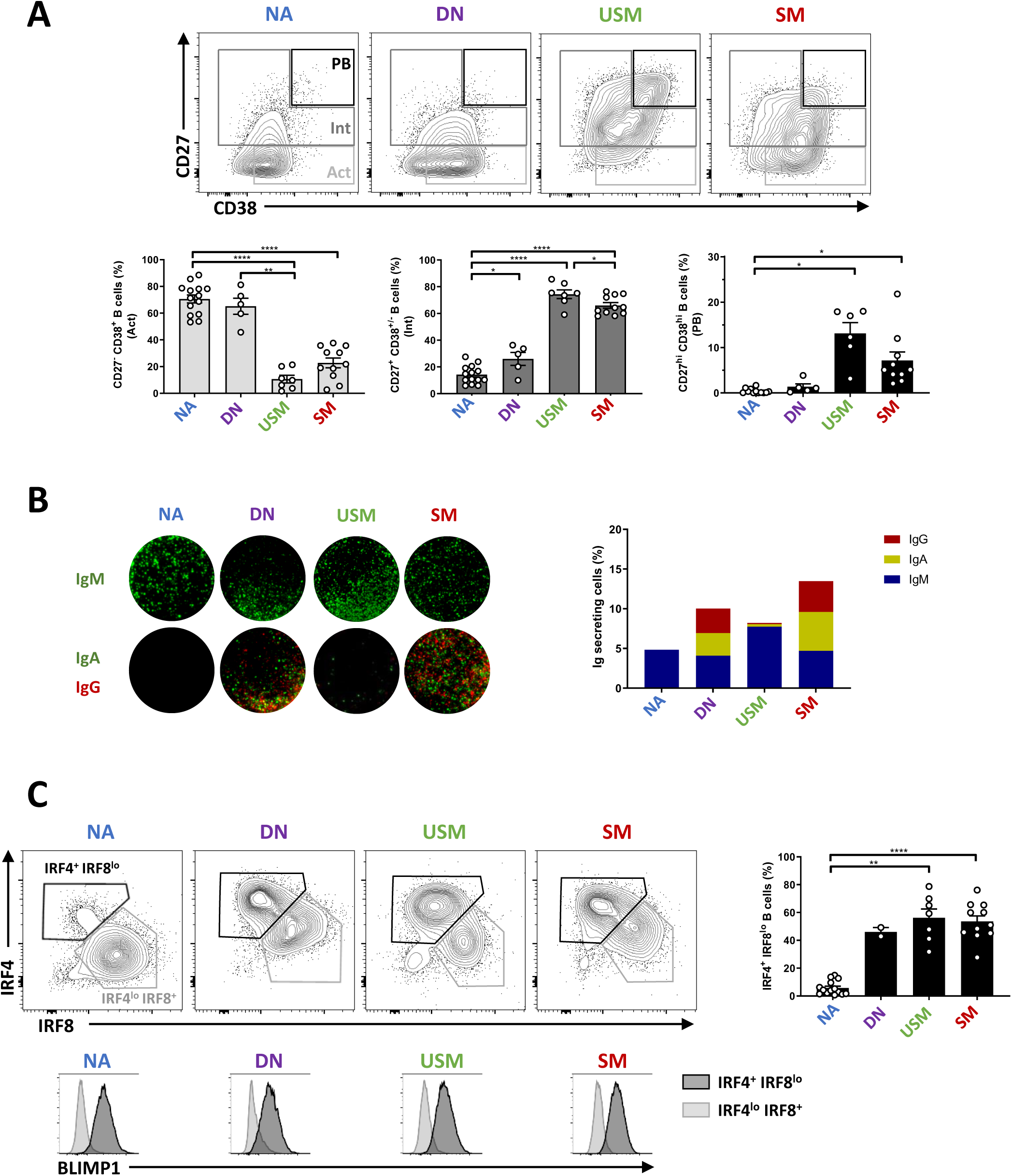
Assessment of the responsiveness of each B-cell subset primed with IL-2, anti-Ig, and CpG. **(A)** Representative flow cytometry plots of CD38 and CD27 expression on CD19^+^ B cells. Three populations, observed after stimulation (priming), are gated: CD27^high^ CD38^high^ B cells (PBs, plasmablasts in black), CD27^+^ CD38^int^ (Int, intermediate B cells in dark grey), CD27^-^ CD38^+^ (Act, activated B cells in light grey). Mean ± SEM percentages (below) of Act, Int and PBs in each B-cell subset. (N= 5 to 14) **(B)** Assessment by ELISPOT of the Ig secretory function in primed B cells. The right plot depicted the percentages of IgM-(blue), IgA-(yellow), IgG-(red) secreting B cells **(C)** Representative flow cytometry plots of IRF4 and IRF8 intracellular staining. Black gates depicted IRF4^+^ IRF8^low^ differentiated B cells and grey gates the IRF4^low^ IRF8^+^ activated B cells. Mean ± SEM percentages (right) of IRF4^+^ IRF8^low^ B cells. Data are representative of N= 14 (NA); N= 5 (DN); N= 7 (USM), and N= 11 (SM) independent experiments. Below are the histogram overlays of the expression of BLIMP1 in IRF4^+^ IRF8^low^ (black) and IRF4^low^ IRF8^+^ (grey) populations. Statistical significance in (A, C) was determined by a one-way paired ANOVA test using Tukey’s correction for multiple comparisons. Error bars indicate mean ± SEM. ** p<0.01, *** p<0.001. NA: Naive B cells, USM: Unswitched memory B cells, SM: Switched memory B cells, DN: Double-negative B cells.

### IL-21 and IFN-α modify the mature B-cell responses and uncover unique innate precursor ability

To investigate the role of IL-21 and IFN-α signaling in mature B-cell differentiation, we assessed the induction of IRF4^high^ IRF8^low^ cells in the primed culture in the presence of IL-21 or IFN-α in the four sorted populations (Figure 2A). A relative heterogeneous rate of cell differentiation existed within the IgD^+^ compartment according to the different donors (Figure 2B). Whereas IFN-α exhibited no significant additive effect to the priming activation on the activation of any mature subset, IL-21 signaling had opposite results between NA and SM cells. Primed SM cells had a 30 % increase in the differentiation rate in the presence of IL-21. On the contrary, IL-21 suppressed the induction of the IRF4^high^ IRF8^low^ population in the NA culture. Those observations were confirmed by ELISPOT at the functional level confirming the loss of ASC coming from the NA population in the presence of IL-21 (Supplementary Figure 1B). IgM/IgD memory and class-switched cells might exhibit a distinct fate after reactivation in mice (28). Thus, we compared the differentiation response to IL-21 in the IgD^+^ population (Figure 2B). In contrast to the effect on IgD^+^ CD27^-^ cells, IL-21 had no additional impact on the differentiation of IgD^+^ CD27^+^ USM cells and did not exhibit a suppressive effect.

**Fig. 2.**
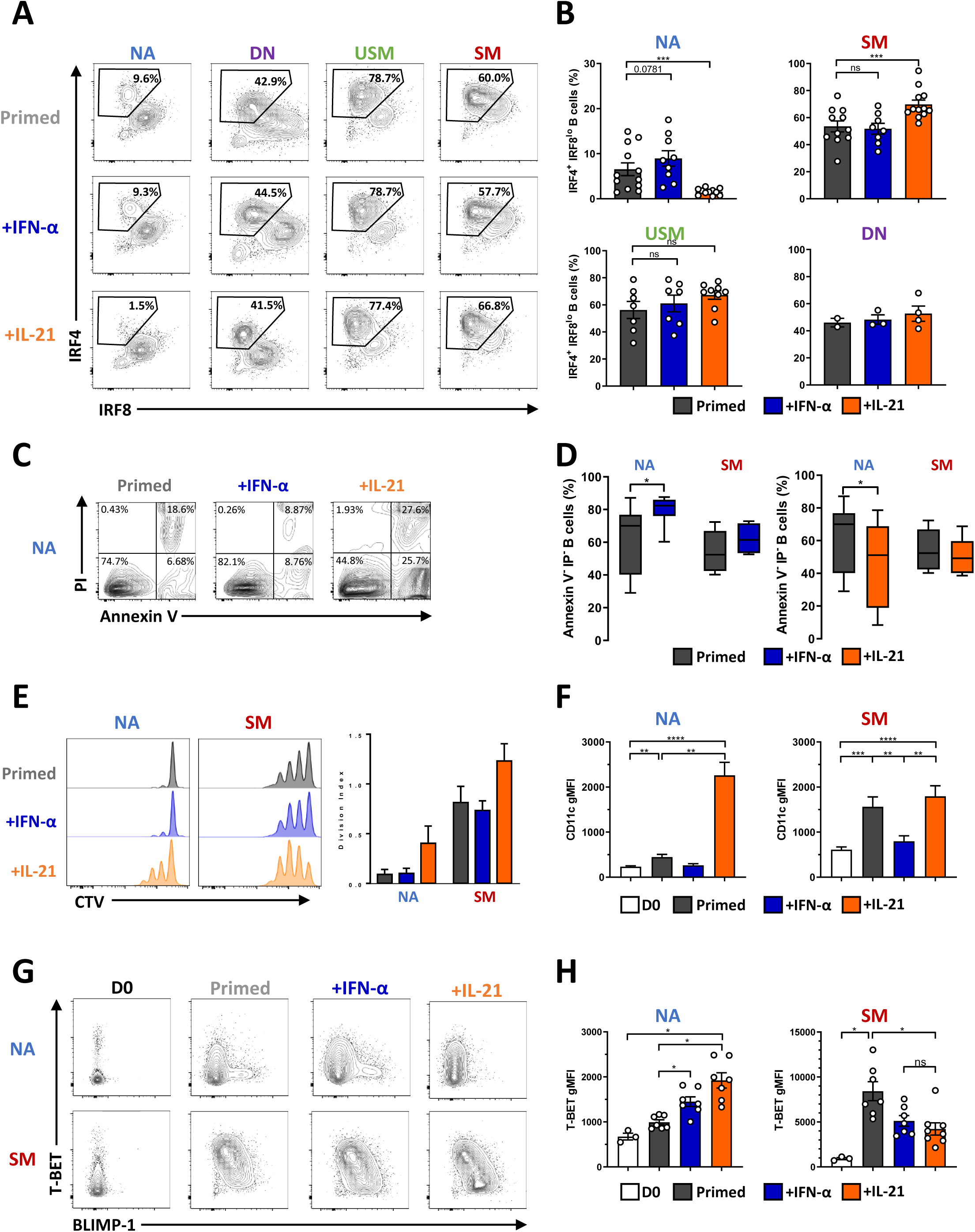
Effects of IL-21 and IFN-α on B-cell responses. **(A)** Representative differentiation profile of the four primed B-cell subsets cultured in the presence or not of IFN-α or IL-21. **(B)** Mean ± SEM of IRF4 IRF8 B cells generated in the different cultures of primed B-cell subsets cultured in the presence or not (dark grey) of IFN-α (blue) or IL-21 (orange). Data are representative of N= 14 (NA), N= 12 (SM), N= 9 (USM) and N= 4 (DN) independent experiments. **(C)** Annexin-V/Propidium Iodure (PI) representative cytometry data of primed NA B cells cultured or not with IFN-α (blue) or IL-21 (orange). **(D)** Mean ± SEM viable Annexin-V^-^ PI^-^ NA and SM primed (dark grey) B cells cultured in the presence or not of IFN-α (blue) or IL-21 (orange). Data are representative of 4 to 7 independent experiments. Statistical significance was determined by paired Wilcoxon paired t-test. **(E)** Assessment of the proliferation of primed NA and SM B cells (left) cultured in the presence or not of IFN-α (blue) or IL-21 (orange). The right panel depicted the mean ± SEM division index (N= 3 independent experiments). **(F)** Mean ± SEM CD11c geometric mean intensity (gMFI) in NA (left) and SM B cells (right) at day 0 (D0, white) and after stimulation (primed) in the presence or not (dark grey) of IFN-α (blue) or IL-21 (orange). N= 8 to 10 independent experiments. **(G)** Representative flow cytometry plots of T-BET and BLIMP-1 expression analyzed by intracellular flow cytometry **(H)** Mean ± SEM geometric mean for T-BET expression on B cells in the different culture conditions. Data are representative of 3 (D0) to 8 independent experiments, and statistical significance was determined by ANOVA test with Tukey’correction. NA: Naive B cells, USM: Unswitched memory B cells, SM: Switched memory B cells, DN: Double-negative B cells.

IL-21 was initially described to deliver both positive and negative signals in a context-dependent manner in NK cells and B cells (29) and more recently on dendritic cells (30). To date, several articles have reported a suppressive effect of IL-21 on normal B cells (31–33). To understand the opposite role of IL-21 on SM- and NA-derived PB differentiation, we examined the effect of IL-21 on the B-cell apoptosis (Figure 2C). The addition of IL-21 to the initial stimulation significantly decreased annexin V^-^ PI^-^ viable NA cells but not viable SM cells. On the contrary, IFN-α significantly promoted NA cell survival, whereas it had no impact on SM cell recovery (Figure 2C and 2D). Multiple rounds of division and differentiation coexist during the reactivation of MBC, balancing *in vivo* between the generation of either GC or plasma cells (34, 35). Furthermore, the functional division of MBCs following reactivation might highly contribute to their regenerative potential. When activated, NA cells remained largely undivided at day 3. In contrast, SM cells proliferated with more than three divisions. IL-21 increased the proliferation of SM cells induced by priming activation, whereas it led NA to enter into the cycle (Figure 2E). We analyzed the apparition of the IRF4^+^ IRF8^low^ PB population in both subsets each day during the culture (Supplementary Figure 1C). There was a notable difference in the velocity of the differentiation process. IL-21 amplified and maintained both rounds of division and differentiation in SM cells initiated at the beginning of the culture. In contrast, cell activation triggered the differentiation of a subset of NA cells with a significant delay compared with that of SM cells. The IL-21-induced proliferation of NA cells might lead *in vitro* to *nutrient exhaustion* and subsequent non-specific apoptosis that may be uncoupled from direct IL-21 signaling. When following the kinetics of the IL-21-dependent apoptosis in NA cells, live imaging revealed a very progressive process that was initiated after 10-15 hours of stimulation, suggesting an activation-induced mechanism (Supplementary Figure 1D).

IL-21 induced T-BET and CD11c expression in anti-IgM/CD40L activated naive B-cell culture (36). As expected, CD11c was strongly induced by IL-21 without any differences between the subsets (Figure 2F). Interestingly, IL-21 decreased the induction of T-BET in activated SM cells together with the induction of BLIMP1, as mutually exclusive events (Figure 2G, and 2H, Supplementary Figure 1E). The addition of IL-21 to activated NA cells led to opposite results, with significant suppression of the BLIMP1^+^ population toward a T-BET-positive phenotype underlining two opposite maturation trajectories.

Altogether, these results demonstrated that IL-21 signaling specified different primary outcomes in B-cell subsets promoting either proliferation and T-BET-activation in NA B cells or both renewal and differentiation in MBC. Moreover, we uncovered a unique NA-derived IgD^+^ CD27^-^ population with delayed ability to differentiate into IgM-producing PB repressed by IL-21.

### IL-21 and IFN-α induce different transcriptional programs in naive and switched memory B cells

To gain insight into the role of IL-21 and IFN-α signaling in activated mature B-cell subsets, we performed RNA sequencing on sorted NA and SM B cells in a resting state and cultured them in the presence of both cytokines. The unsupervised approach showed an accurate clustering of B-cell subsets according to their origin and *stimuli* (Figure 3A). Principal component analysis 1, representing 54% of the total variance, is correlated with the culture-activation conditions, whereas PC2 (15% of the variance) separated the two major B-cell subsets (Supplementary Figure 2A). We treated the loadings of the principal components onto each of the genes as expression data to run pathway analysis using the PAGE algorithm on multiple GO pathways and hallmark.MsigDB databases (37) (Figure 3B). As anticipated, PC1, representing the culture activation, was strongly associated with the mitotic process, cell cycle, and DNA replication, whereas PC2 correlated with the reticulum process, mRNA translation, and protein production. We then analyzed genes differentially expressed in NA cells in the presence of IL-21 compared to the priming stimulation (adjusted p value = 0.05, fold change = 1.5). Although the sensitivity of our dataset was modest, we observed that differentially expressed genes (DEGs) reflected the principal cellular outcomes in cultures between the different conditions (Figure 3C and Supplementary Table 3). Among the 124 genes down-regulated by IL-21, we found genes involved in Ig transcription and protein production (*XBP1*, *BHLHA15*, *CAV1*, *TNFSF8*, *FKBP11*, *MZB1*, and *IGVK*) were both key elements of B-cell differentiation into ASCs (Figure 3C). In contrast, the presence of IL-21 strongly up-regulated genes involved in the control of B-cell activation and proliferation of *AICDA*, *TOX*, *TOX2*, *NR4A1*, *ZBTB32* and, *ITGAX* (CD11c) gene expression. GO enrichment analysis highlighted the up-regulation of metabolic activity in IL-21 stimulated NA cells with the enrichment of kinase, transferase, fatty-acyl-CoA, and lipid metabolic processes (Supplementary Figure 2B). Similar observations were made when we compared NA cells stimulated either with IL-21 or IFN-α (Figure 3D). IFN-α-stimulated NA B cells exhibited induction of *PRDM1*, *CD38*, and *XBP1* involved in PB differentiation and Ig transcription whereas the genes induced in the presence of IL-21 (*AICDA*, *TOX*, *BHLHE40*, *CD276*, *PBX4*, and *IKZF4*) suggested a more GC-like pathway (38, 39) (Figure 3D). As expected, the IFN-α signaling is strongly emphasized in NA cells by the up-regulation of an important set of type I IFN-responsive genes (*IRF7*, *MX1*, *MX2*, *IFI44*, *IFI35*, *ISG15*, *OAS3*, *IFI6*, and *IFI16*) (Supplementary Figure 2C).

**Fig. 3.**
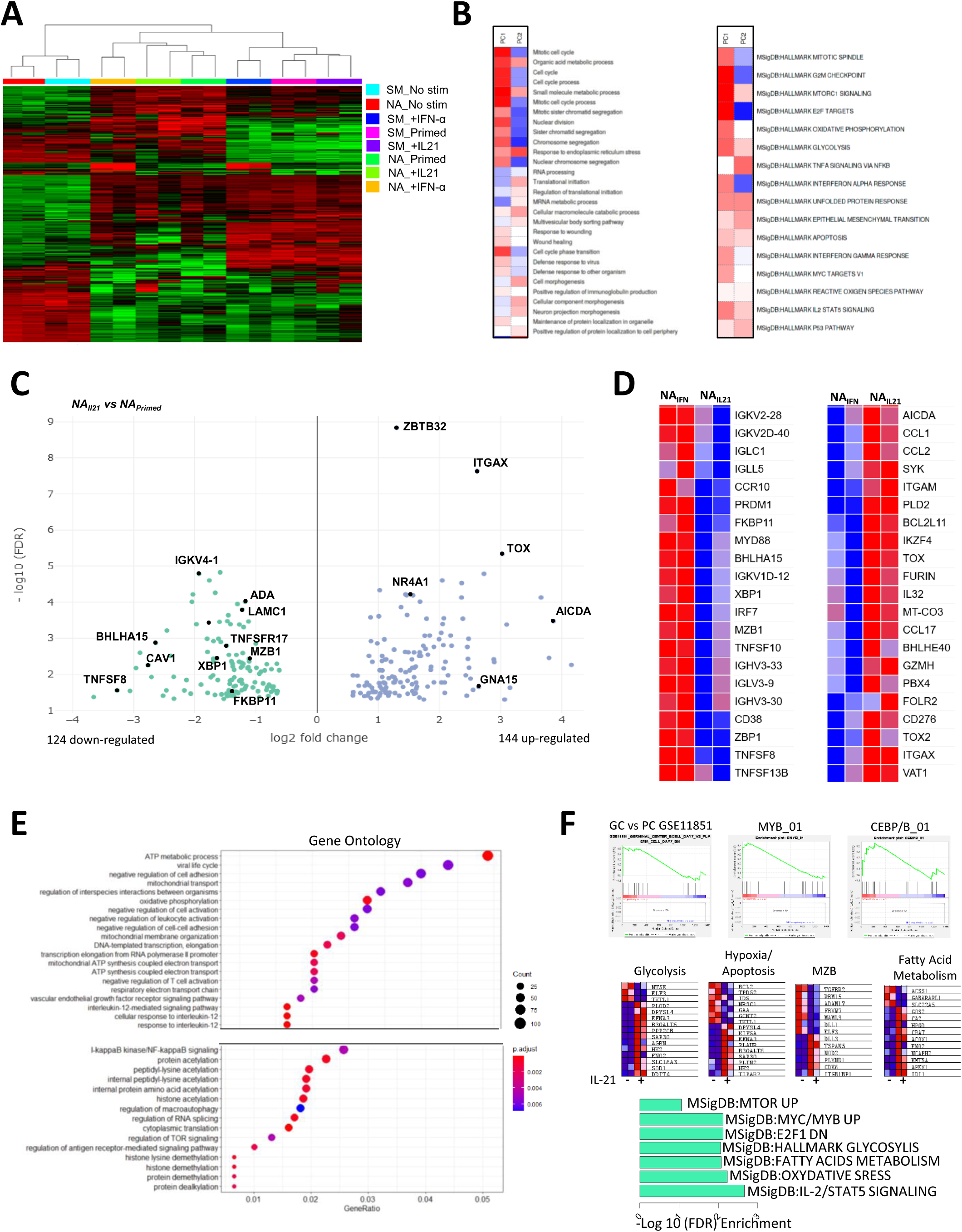
Transcriptional programs induced by IFN-α and IL-21 in naive and switched memory primed B cells. **(A)** Unsupervised hierarchical clustering of gene expression in NA and SM cells at resting state (No stim), after stimulation in the presence or not of IFN-α or IL-21. **(B)** Heatmap of the principal component 1 and principal component 2 associated pathway from the PGSEA analysis. **(C)** Volcano plot of DEGs between NA primed B cells (NA_Primed_) and NA cells primed in the presence of IL-21 (NA_IL21_). (**D**) Heatmap of important genes differentially expressed between NA primed B cells in the presence of IFN-α (NA_IFN_) *versus* IL-21 (NA_IL21_). (**E**) Plot of Gene Ontology (GO) pathway enrichment of the genes only upregulated by the addition of IL-21 in NA primed cells. (**F**) Gene set enrichment analysis (GSEA) (upper), genes in major pathways (middle), and MSigDB pathway enrichment of the IL-21 signature in NA cells (bottom). NA: Naive B cells, SM: Switched memory B cells.

Since the IL-21 signaling was initiated at the beginning of the culture, we hypothesized that important variations in gene expression occurred during the 3 days of stimulation and could be reflected by assessing DEGs during activation with or without IL-21 related to unstimulated NA B cells (Supplementary Figure 2D, Supplementary Table 3). The overlapping profile of both simulations revealed a core activation signature that reproduce changes in gene expression mediated by the priming stimulation. A set of 819 up-regulated and 795 down-regulated genes were modulated only in the presence of IL-21. In agreement with the increase of mortality observed in the culture, *CFLAR*, *DDIT4*, *BCL2L11* (BIM), *BNIP3*, and *PAK2* apoptotic genes were up-regulated and the anti-apoptotic protein BCL2 was down-regulated exclusively in the presence of IL-21 (Supplementary Table 2). Gene ontology, KEGG, and pathway gene set enrichment analysis (GSEA) was used to characterize the IL-21 signature in NA B cells. Metabolic processes appeared as recurrent up-regulated pathway and included the increase of oxidative phosphorylation, fatty acid metabolism and glycolysis (Figure 3E and 3F, Supplementary Figure 2E). These pathways seem to be associated with an enrichment of mTOR activity. The IL-21 signature in activated NA cells was further characterized by an enrichment of a C/EBPβ and MYB-dependent-genes network over the MZB-related genes (*DLL1*, *ADAM17*, *MALM3*, *FBXW7*, *RBM15*, and *TGFBR2*) (Figure 3F) (40–42).

Finally, we compared the effects of IL-21 signaling after activation between the NA and SM cell subsets resulting in 1227 up-regulated and 1166 down-regulated genes (Supplementary Figure 2F and Supplementary Table 4). These DEGs corroborated the interpretation of the different fates mediated by IL-21 on the two B-cell subsets. IL-21-stimulated NA cells harbored a transcriptional program close to pre-GC cells with the up-regulation of *BCL6*, *BACH2*, *SPIB*, *IRF8*, and *TCL1* whereas IL-21 induced a strong differentiation of SM cells into PB illustrated by the increase of *PRDM1*, *IRF4*, and *XBP1* and the down-regulation of B-cell lineage associated genes, including *CD22*, *CD72*, *MS4A1*, *CD19*, and *PAX5*) (Supplementary Figure 2G).

Altogether, our transcriptional analyses argued for the down-regulation by IL-21 of PB differentiation in the NA B-cell subset that is conserved in SM cells. Moreover, these data established B-cell-intrinsic IL-21 signaling in primed NA B cells with a high metabolic signature led to proliferation and activation-induced apoptosis.

### CD45RB, CD1c, and unique homing receptor profile identify a circulating CD27^-^ marginal zone B-cell-related progenitor population

We uncovered the existence of a B-cell population within the IgD^+^ CD27^-^ NA compartment that rapidly differentiated upon BCR and TLR signaling but had opposite responses between IL-21 and IFN-α. This identity coincided with the down-regulation of some genes involved in MZB differentiation. We thus hypothesized the existence of a MZB-related progenitor cell population displaying multi-potential properties. To explore this idea, we conducted an extensive phenotypic analysis using mass cytometry on circulating peripheral blood (Figure 4). The CD45RB isoform detected by the Ab clone MEM-55 was reported as an accurate marker to identify not only CD27^-^ and CD27^+^ MBCs (43, 44) but also MZB cell lineage subsets (45, 46). We thus used a routine panel of 35 markers including anti-CD45RB (MEM-55) Ab and the intracellular detection of T-BET on whole blood from seven different donors (Figure 4A). The ViSNE algorithm was run on the concatenated CD19^+^ B-cell data to visualize B-cell populations projected onto a two-dimensional map (Figure 4A). CD27 expression distinguished two distinct areas on the ViSNE cell map, and IgD^+^ cells represented the majority of mature B cells. We then performed unsupervised hierarchical clustering using FlowSOM resulting in 15 meta clusters overlaid on the ViSNe plot with various IgD, CD27, and CD45RB density expressions (47) (Figure 4B). Among them, cluster 12 (C12) represents a unique population of B cells among the NA CD27^-^ population expressing IgD and high density of CD45RB. We then applied on the t-distributed stochastic neighbor embedding (t-SNE) the overlay of the manually gated IgD^+^ CD27^-^ CD45RB^+^ (NARB^+^) population together with the most currently known B-cell subsets, including the two newly described B-cell populations poised to differentiate into PB and expanded in autoimmune patients referred to as activated naive (aNav) and DN2 cells (Supplementary Figure 3) (21, 48). The NARB^+^ population does not express T-BET or CD11c as described for the aNav and DN2 cell populations that appeared to cluster together in C3, confirming the previously described relationship between them (Figure 4C). The NARB^+^ cells were characterized by a high amount of the B-cell follicle homing receptors CXCR5, CCR7, and CCR6 but low expression of CXCR3. This profile contrasted with USM and SM cells that exhibited the up-regulation of CXCR3 expression and down-regulation of CCR6 expression. We further realized an extensive identification of this population with additional markers in other donors. The NARB^+^ population represented 4.5 ± 1.8% of total B cells with a relatively stable representation, contrasting the importance of donors’ variability of human CD27^+^ MBCs (Figure 4D). NARB^+^ presented an intermediate profile between NARB^-^ B cells and USM B cells (IgD^+^CD27^+^) and can be identified as IgM^+^ IgD^high^ CD45RB^+^ CD27 MTG^int^ CD38^-^ CD23^low^ CD21^high^ CD1c^+^ CD80^low^ PDL2^-^ B cells (Figure 4E). The circulating NARB^+^ population shared a common phenotypic identity (expression of IgM and CD1c) with IgD^+^ IgM^+^ CD27^+^ MZ-like B cells but with a very distinctive homing (CXCR4, CXCR5, CCR6, and CCR7) and signaling-associated molecules (CD21, CD23, CD22, CD5, TACI, CD80, PDL2, and CD32) profile.

**Fig. 4.**
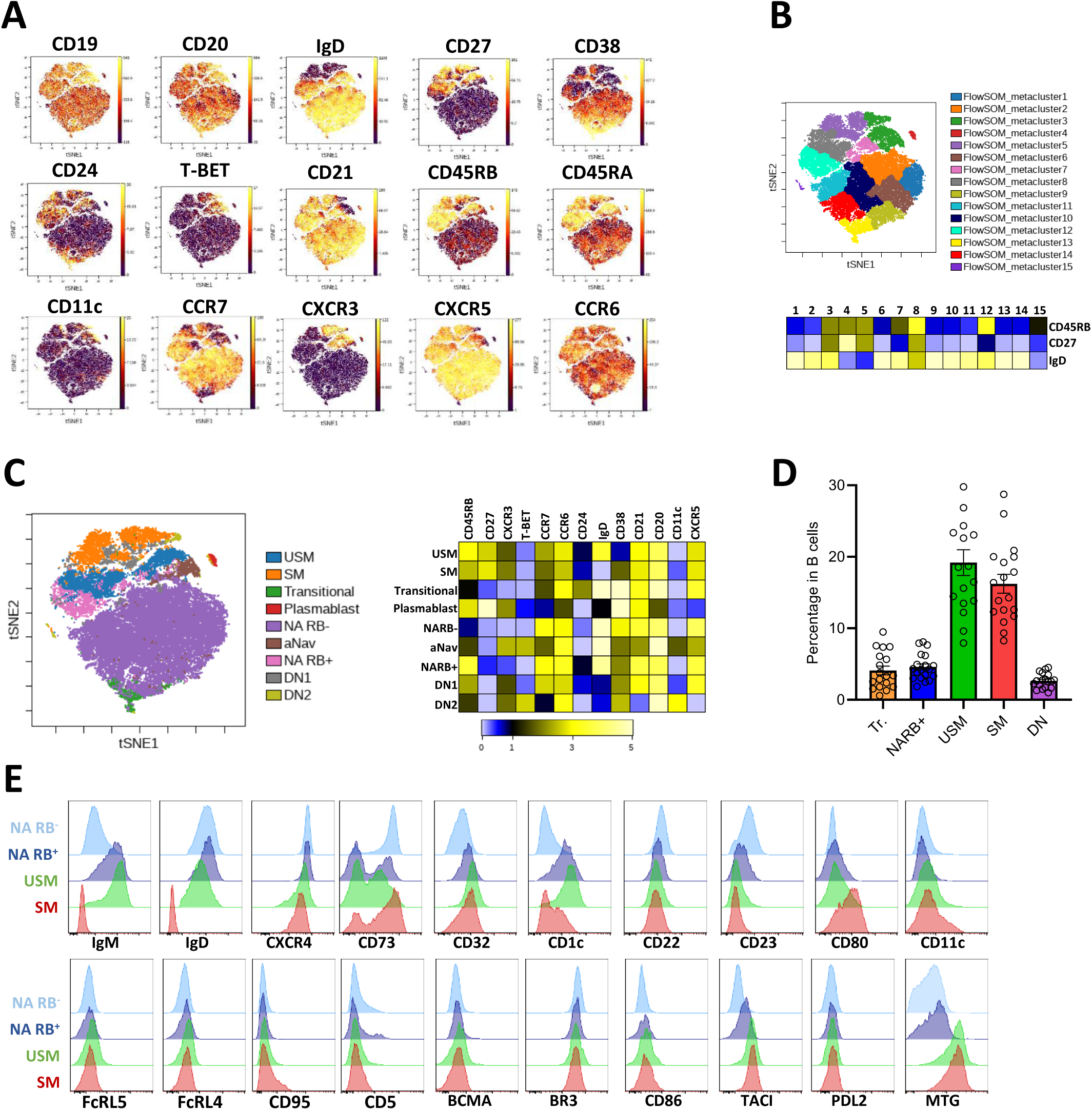
Phenotypic characterization of IgD^+^ CD27^-^ CD45RB^+^ cells using CyTOF and Flow cytometry. **(A)** VisNE analysis of mass cytometry data performed on B cells from whole blood of healthy control (concatenate files from 5 donors). **(B)** FlowSOM clustering with the representation of the 15 meta-clusters on the t-SNE map and heatmap of expression density of IgD, CD27, and CD45RB markers (below). **(C)** Manual gating of the major peripheral B-cell subsets loaded on the t-SNE plots and heatmap of expression density of selected markers (below). **(D)** Mean ± SEM of the percentage of transitional (Tr.), NARB^+,^ USM, SM, and DN subsets in B cells. Data are from 18 independent donors. **(E)** The extended phenotype of NARB^+^ cells compared to NARB^-^, USM and SM B cells performed by flow cytometry. NA: Naive B cells, NARB^+/-^: CD45RB^+/-^ naive B cells, aNav: activated naive B cells, USM: Unswitched memory B cells, SM: Switched memory B cells, DN: Double-negative B cells.

### NARB^+^ cells are uncommitted precursors of innate plasmablasts

According to these data, we thus hypothesized that this progenitor population might represent the previous B-cell subset among the NA cells that exhibit rapid differentiation upon reactivation. To confirm this hypothesis, we sorted NARB^+^ and NARB^-^ cells to evaluate their response in the priming culture. The induction of the IRF4^+^ IRF8^low^ BLIMP1^+^ PBs was only observed in the NARB^+^ cells culture (Figure 5A and 5B) identifying NARB^+^ as the precursor population of IgM^+^ IgD^+^ CD27^+^ cells and IgM^+^ PBs. We followed the apoptosis of sorted NARB^+^ and NARB^-^ cells by live-cell imaging in the primed culture with or without IL-21. NARB^+^ cells conserved functional characteristics of Ag-inexperienced NA cells in response to IL-21 exhibiting rapid activation-cell death with similar kinetics (Figure 5C).

**Fig. 5.**
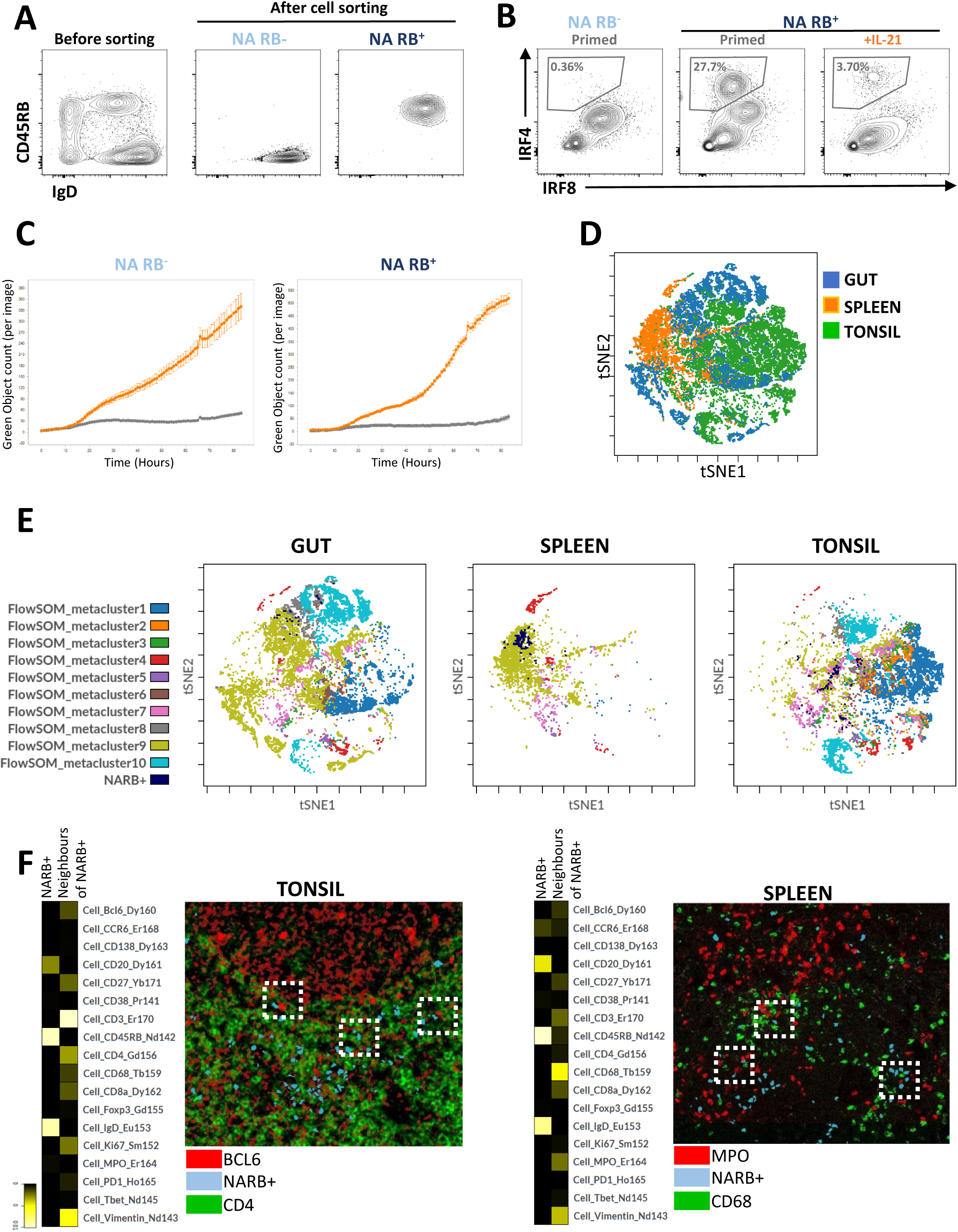
NARB^+^ cells are non-committed precursors of innate plasmablasts and localized in secondary lymphoid organs. **(A)** NARB^+^ B-cell sorting purity. **(B)** Representative FACS plots of IRF4 and IRF8 intracellular staining in NARB^-^ and NARB^+^ primed in the presence or not of IL-21. Black gates indicate IRF4^+^ IRF8^low^ differentiated B cells. **(C)** Kinetics of apoptotic annexin-V^+^ B cells in NARB^-^ and NARB^+^ stimulated in the presence (orange line) or not (grey line) of IL-21, evaluated by live-cell imaging. **(D)** The t-SNE plot was performed on individual cell files (FCS) obtained from imaging mass cytometry data from spleen, gut, and tonsil tissue sections. **(E)** FlowSOM clusters were loaded the t-SNE plots in gut, spleen, and tonsil tissues together with NARB^+^ cells previously manually gated. **(F)** Heatmap view of the neighborhood analysis of NARB^+^ cells in the tonsil and spleen. Representative images of BCL6^+^, NARB^+^ and CD4^+^ cells in the tonsil, and MPO^+^, CD68^+^, and NARB^+^ in the spleen. The dashed square lines identified cell-cell interactions. All data were generated with histoCAT tool and Cytobank.

We then explored whether these cells could be found in secondary lymphoid organs at steady state. To address this question, we performed imaging mass cytometry on tonsil, spleen, and gut-associated lymphoid tissue sections (Supplementary Figure 4A). We visualized and analyzed data using the open-source platform histoCAT, which enables highly multiplexed, quantitative, and detailed analysis of cell phenotypes, microenvironment interactions, and tissue architecture. The workflow of histoCAT consists of overlaying the segmentation masks previously generated by CellProfiler to extract single-cell level information and processing this information into FCS files. We analyzed FCS files analyzed using the t-SNE algorithm that revealed specific cell distribution according to tissue origin (Figure 5D). Unsupervised hierarchical clustering using the FlowSOM algorithm was performed to cluster 10 major groups of cells in the different tissues (Figure 5E). Although the expression levels of the different markers may significantly vary between them and across the tissues, the heat map view of the FlowSOM clusters identified clusters 1 and 2 were mostly associated with T cells; cluster 3 with macrophages; cluster 4 with neutrophils, and clusters 8, 9, and 10 with B cells. Clusters 5, 6 and 7 had a more ambivalent profile. Finally, Ki67^+^ proliferative B cells identified the GCs in tonsil and the gut but not in the spleen sections analyzed here (Supplementary Figure 4B). We confirmed that the clusters overlapped with the main cell populations identified by manual gating. NARB^+^ B cells were detected belonging mostly to cluster 9 (Figure 5E and Supplementary Figure 4C). NARB^+^ cells were localized outside the GC mostly in the mantle zone as previously described for IgD^+^ cells. Using neighborhood analysis with histoCAT, NARB^+^ cells were found in very close interaction with CD3^+^CD4^+^ BCL6^+^ T cells localized at the border of the GCs in the tonsil and in the gut-associated tissues. Moreover, NARB^+^ appeared to be close to Ki67^+^ proliferative CD3^+^ T cells that might suggest antigen presentation or activation process. However, the cell interaction map of NARB^+^ differed in the spleen where NARB^+^ were mostly in interaction with innate cells like with CD68^+^ macrophages and MPO^+^ neutrophils (Figure 5F and Supplementary Figure 4D). These data suggested circulating NARB^+^ cells had equivalence in secondary lymphoid tissues that were not restricted to the spleen and might participate in the immune response.

### NARB^+^ cells are in a pre-activated state with unique pSTAT3 and pSTAT1 pattern

IL-21 signaling *via* downstream STATs phosphorylations specified the outcome of naive and memory B cells (12, 19). To shed light on the mechanisms underlying the specific effects of IL-21 on the NARB^+^ population, we compared the early signaling events triggered by IL-21 in combination with the primed stimulation in the different subsets. IL-21 signals through the receptor pair IL-21R/γc, which leads to induction of STAT3 and STAT1 and to a lesser extent the STAT5 and PI3K/AKT pathways. In the priming stimulation, the STAT3 activation showed a similar dynamic of phosphorylation between the different subsets with a peak after 5 min and then a decrease. The STAT1 phosphorylation in USM, SM, and NARB^+^ cells also had a peak of activation after 5 min, however it continued to gradually increase between 30 and 90 min. This increase was not observed for NARB^-^ cells (Figure 6A). We analyzed the phosphorylation levels of Y705-STAT3 and Y701-STAT1 in the different B-cell subsets after 30 min of activation with or without IL-21 (Figure 6B). In the presence of IL-21, NARB^+^ cells presented a dual profile characterized by an important increase of pSTAT3 as observed in Ag-inexperienced NARB^-^ cells and a decrease of pSTAT1 as observed in MBC (USM and SM). In addition to STATs activation, the IL-21 proliferative effect was dependent on the PI3K/AKT pathway (49). We thus examined S473-AKT phosphorylation in the different subsets following activation (Figure 6C). We did not detect a strong additional effect of the addition of IL-21 on the AKT phosphorylation in any of the subsets. However, as observed for STATs phosphorylation, a strong difference existed between the magnitude of the pAKT response following activation between NARB^-^ and NARB^+^ cells. Altogether, these data showed that NARB^+^ cells exhibited a specific IL-21-dependent pSTAT1 and pSTAT3 activation pattern. Moreover, NARB^+^ cells have an intracellular-phosphorylation signature close to Ag-experienced cells suggesting their previous *in vivo* activation.

**Fig. 6.**
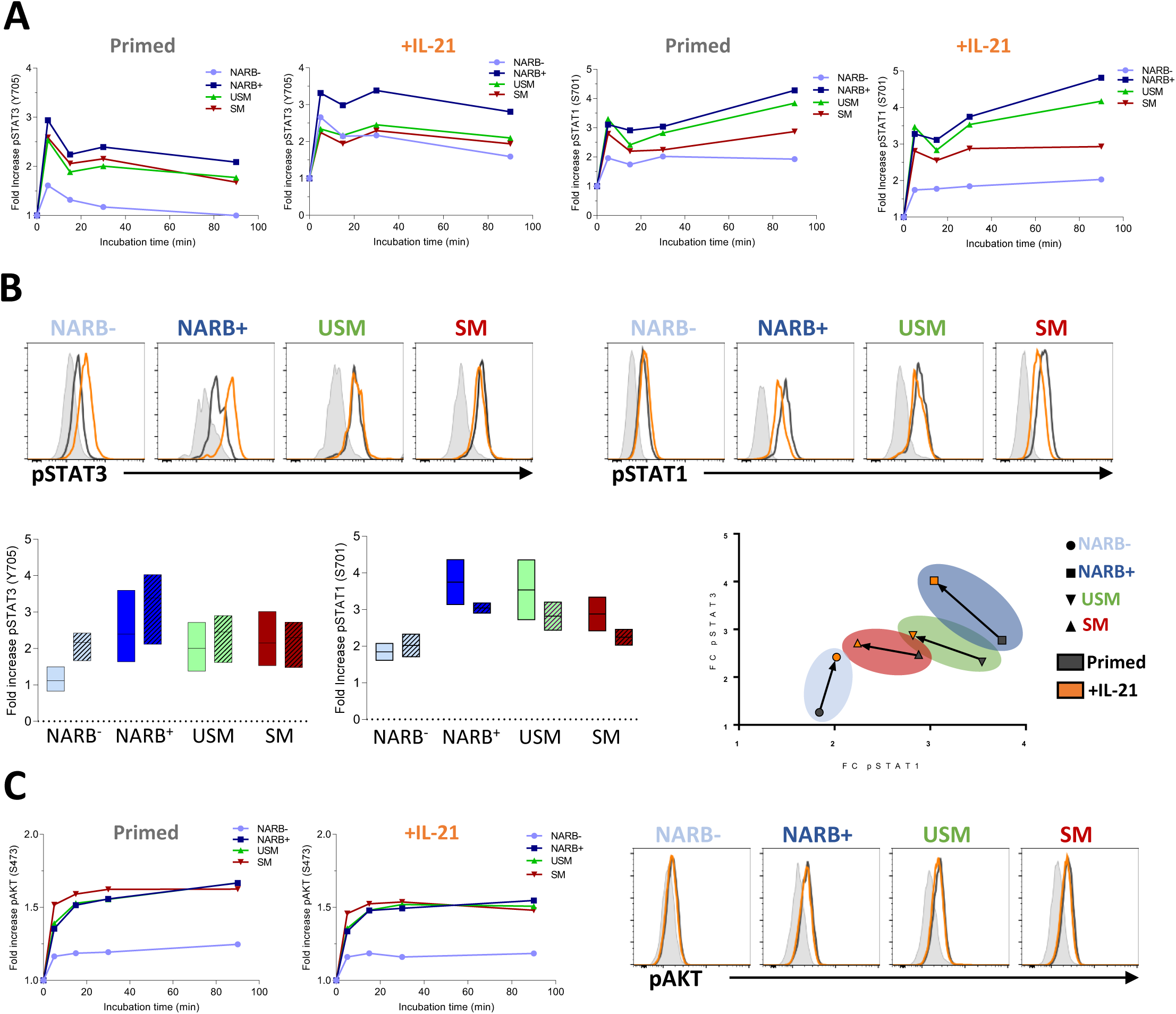
NARB^+^ cells present a unique phosphorylated STATs profile. **(A)** Fold increase of pSTAT3 (Y705) (left) and pSTAT1 (S701) (right) in primed NARB^-^, NARB^+^, USM, and SM subsets cultured in the presence or not of IL-21 at 5, 10, 30 and 90 min. **(B)** Representative flow cytometry plots of pSTAT3 (left) and pSTAT1 (right) in primed NARB^+^, NARB^-^, USM, and SM B-cell subsets (grey line) cultured in the presence (orange line) or not of IL-21. Fold increase of pSTAT3 and pSTAT1 compared to the primed stimulation (N= 3). **(C)** Fold increase of pAKT (S473) in primed NARB^-^, NARB^+^, USM et SM B-cell subsets (left) cultured in the presence (middle) or not of IL-21 at 5-, 10-, 30- and 90-min. Representative flow cytometry plots of pAKT (right) in primed NARB^-^, NARB^+^, USM et SM subsets (grey line) cultured in the presence (orange line) or not of IL-21 (N= 3). NA: Naive B cells, NARB^+/-^: CD45RB^+/-^ naive B cells, aNav: activated naive B cells, USM: Unswitched memory B cells, SM: Switched memory B cells.

### IFN-α but not apoptosis inhibition counteracted IL-21 mediated suppression of NARB^+^ differentiation

*In vitro* activation-induced apoptosis may result from an absence of optimum survival signals potentially available *in vivo*. Quantitative RT-PCR performed on four independent experiments showed that IL-21 down-regulated *BCL2* mRNA together with several other B-cell survival and pro-differentiating factors (*BAFF*, *IL6*, *IL6R*, and *IL10*) in NA cells, confirming the observations obtained from RNA sequencing (Figure 7A). NARB^+^ produced twice as much IL-6 as NARB^-^ at steady state (Figure 7B). After *in vitro* activation, the IL-6 production in NA cells was reduced in the presence of IL-21 whereas IL-10 appeared to be unmodified (Figure 7C and 7D). We thus tested whether supplementation of survival factors in the culture might rescue NA cells from IL-21-dependent apoptosis and restore NARB^+^ differentiation. The addition of IL-6 and IL-10 to the IL-21 primed condition at different doses neither improved the viability of NA cells nor the induction of IRF4^+^ PB in the culture (Figure 7E). The CD40 signaling is described as a GC key pro-survival factor and is a hallmark of the provision of T-cell help to B cells. Accordingly, the addition of a CD40L multimer (Figure 7F and 7G) or BAFF (data not shown) in culture inhibited IL-21-induced apoptosis but did not induce NARB^+^ cells to differentiate, suggesting a subset-intrinsic control of PB differentiation. We then examined whether the biochemical blocking of caspase-induced apoptosis could restore NARB^+^ PB differentiation. We pretreated NA cells with the pan-caspase inhibitor Z-VAD before their activation in the presence of IL-21. As expected, we obtained a full recovery of cell viability in culture, but the PB differentiation was not restored decoupling both mechanisms (Figure 7F and 7G). Lastly, as a test of whether innate signals could overcome IL-21-induced inhibition of NARB^+^ cell differentiation, we analyzed the effect of adding IFN-α to the IL-21 primed condition (Figure 7H). Although apoptosis of NA cells was only partially corrected, IFN-α counteracted IL-21 signaling in NARB^+^ cells and re-established PB differentiation.

**Fig. 7.**
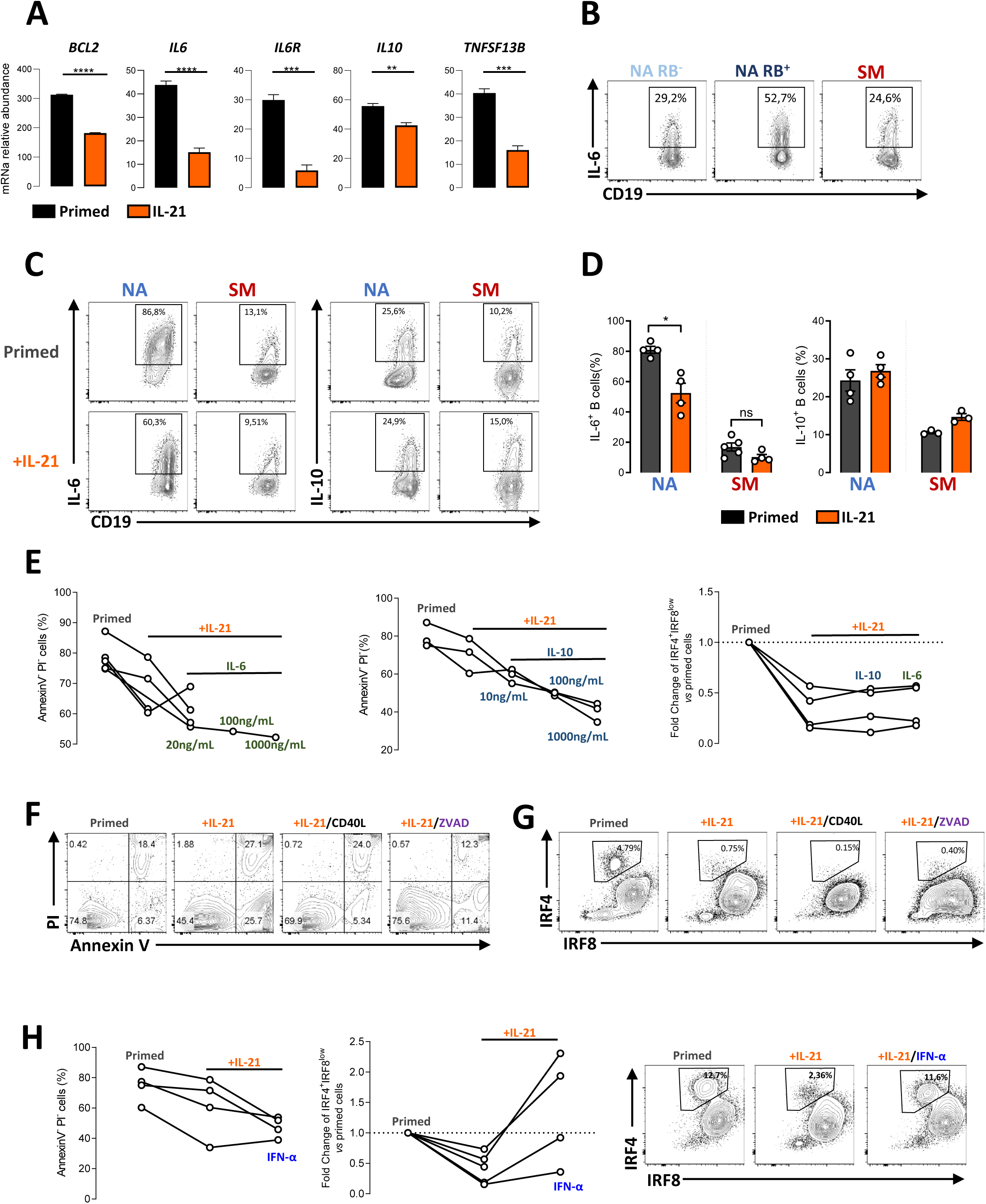
NARB^+^ differentiation was not restored by the inhibition of apoptosis but with innate signals. **(A)** Real-time PCR of *BCL2*, *IL6*, *IL6R*, *IL10* and *TNFSF13B* gene expression in primed NA cells in the presence (orange) or not (black) of IL-21. **(B)** Representative flow cytometry plots of IL-6^+^ cells in NARB^-^, NARB^+^ and SM B cells at steady state. **(C)** Representative flow cytometry plots of IL-6^+^ and IL-10^+^ cells in NARB^-^, NARB^+^, and SM B-cell primed subsets cultured in the presence or not of IL-21. **(D)** Histograms of IL-6^+^ and IL-10^+^ B cells in primed-NA and primed-SM (black) cultured or not with IL-21 (orange). **(E)** Percentage of annexin V^-^ PI^-^ viable cells in primed NA B cells cultured in the presence or not of IL-21, with or without addition of IL-6 (left) or IL-10 (middle). Right plot depicted the fold increase of IRF4^+^IRF8^low^ B cells compared to the primed stimulation. **(F)** Representative plot of AnnexinV/PI staining of NA cells primed cultured in the presence of IL-21, CD40 stimulation (CD40L) or preincubated with the pan-caspase inhibitor (ZVAD) before the culture in the presence of IL-21 **(G)** Representative flow cytometry plots of IRF4 and IRF8 intracellular expression in primed NA cells cultured in the presence or not of IL-21, CD40 stimulation or pan-caspase inhibitor (ZVAD) **(H)** Percentage of AnnexinV^-^ PI^-^ viable cells in primed NA B cells cultured in the presence or not of IL-21 with or without the addition of IFN-α. Middle panel depicted the fold increase of IRF4^+^ IRF8^low^ B cells compared to the primed stimulation. The right panel depicted representative flow cytometry plots of. NA: Naive B cells, NARB^+/-^: CD45RB^+/-^ naive B cells, SM: Switched memory B cells.

Taken together, these data support that NARB^+^ cells not only preferentially followed an IFN-α-dependent pathway of differentiation independently of apoptosis but also kept the ability to proliferate in response to IL-21 and T-cell signals.

## DISCUSSION

In this work, we analyzed the effects of IL-21 and IFN-α signals on different subsets of human activated B cells. We used an *in vitro* B-cell activation model involving BCR and TLR stimulation in presence of IL-2 during a relatively short-time period (referred to as priming cells) (23). The requirement of permanent BCR engagement with an Ag for MBCs to survive or be reactivated is a major debate (50–52), and activation by TLR agonists or bystander T cells help have been suggested to also ensure the longevity of MBC in an Ag-independent process (53). The priming model presented here was an attempt to integrate cognate B-cell stimulation signal 1 and TLR innate signal 2 without direct provision of T-cell help. We demonstrated that this stimulation was sufficient to capture the early events of the activation of memory and naive B cells and then address the ability of MBC to rapidly differentiate into PB. We thus efficiently tracked the IRF4^+^ IRF8^low^ PB and pre-PB subsets with Ab secreting ability regardless of CD27 and CD38 expression. However, this model has the inherent limitations of *in vitro* studies using a limited competitive cells environment and fixed levels of BCR stimulation. In this regard, the affinity and valence of the BCR in the regulation of primary events conducting human B cells toward follicular or EF pathways remain to be addressed (54).

Our data showed that the IFN-α and primed stimulation have a redundant role in the differentiation of MBC subsets into PB. In contrast, IL-21 signaling had a unique intrinsic effect influencing B-cell fate. This specific IL-21-dependent B-cell response led us to uncover the existence of an innate-like mature IgD^+^ CD27^-^ CD45RB^+^ CD1c^+^ CD10^-^ B-cell subset harboring progenitor abilities. This subset presented a versatile “multilineage” differentiation potential, either being able to differentiate into IgM-producing PB upon innate signals or becoming proliferative and apoptotic in the presence of IL-21. NARB^+^ circulating B cells exhibited the expression of follicular homing chemokine receptors and had equivalents in multiple secondary lymphoid tissues which strengthen its progenitor nature. However, NARB^+^ cells differed from transitional B cells notably by the low expression of CD38, the intermediate MTG extrusion, and the absence of CD10 expression. An equivalent subset was previously identified in the spleen expressing the CD45RB glycosylated epitope MEM-55 and up-regulating CD27 expression in response to NOTCH2-ligand classifying those subsets as marginal zone precursor cells (MZP) (17, 55). The CD45RB isoform was initially associated with MBC subsets in humans with a high expression on switched CD27^+^ B cells, PBs, and IgD^+^ IgM^+^ CD27^+^ USM B cells (45). More recently, extensive phenotypic and functional analysis of human peripheral B-cell subsets through mass cytometry has refined the CD45RB^+^ CD27^-^ population as an early memory B-cell subset based on the level of mutational burden, and their unique metabolic and immune signaling activities, however, with no distinction between isotype status (43). In a recent elegant study, the Spencer’ group identified CD45RB^+^ T2 cells as progenitors of human marginal zone B cells, suggesting a very close developmental link with the CD45RB^+^ naive population described in this paper (62). CD45RB^+^ IgD^+^ CD27^-^ (NARB^+^) cells shared different molecular and functional attributes with both naive and memory B cells but with unique IL-21 responsiveness. NARB^+^ cells were able to differentiate upon innate signals toward IgM-producing cells as observed for the IgD^+^ CD27^+^ (USM) subset. Whereas IL-21 signaling had nearly no impact on the priming activation of USM cells, NARB^+^ cell differentiation is suppressed toward activation-induced apoptosis. The reduction of cell viability in the presence of IL-21 is not restrained to NARB^+^ cells but is a feature of the whole naive compartment. Intracellular STAT and PI3K signaling analysis of the different populations following stimulation demonstrated that the NARB^+^ subset exists in a pre-activated state. This finding challenged the belonging of this subset to the Ag-inexperienced compartment. Although our data could not trace the trajectory of NARB^+^ differentiation, the phenotypic observations seemed to place this subset in a developmental continuum from naive B cells to MBC. However, another complementary point of view will be to see NARB^+^ cells as circulating innate progenitors of the MZ response. Thus, NARB^+^ cells could be assimilated to a MZP subset, however, the IL-21 typical response observed *in vitro* suggested that NARB^+^ cells are not fully committed to MZ differentiation and may keep the ability to follow the follicular response in a specific environment. The addition of CD40L signaling to the IL-21 activation cocktail is in favor of this idea, with almost entire cell recovery and entry into the cell cycle. At this stage, we could only speculate on this putative cell pivotal state but its existence in secondary lymphoid organs suggested active participation of the immune response. Furthermore, our mass cell imaging analyses revealed that NARB^+^ cells seem interact more specifically with innate cells (neutrophils and macrophages) in the spleen, and with pre-Tfh in tonsils and in the gut. These data supported that NARB^+^ cells have specific cell-cell interactions according to their tissue localization that might shape their ultimate function. CD27^dim^ MBCs were recently described as enriched in infants and might not only represent a reservoir for CD27^high^ memory effectors but also might enter the GC reaction (56). The NARB^+^ cells described in this report presented similar functional attributes, suggesting a developmental relationship.

IL-21 production is one of the hallmarks of Tfh identity and is related to the control of the GC B-cell response (39). However, the IL-21 activity area is not restrained to the follicular response, and BCL6^+^ PD1^low^ helper T cells located at the T-B border could contribute to the EF response. There was a defect of EF plasma cells in IL-21R and IL-21 knockout mice. It was further proposed that IL-21 could act at the stage of T–B interaction to accelerate B-cell activation before their differentiation into GC cells or to promote EF PB differentiation (6). More recently two Tfh effector stages were described according to their spatial positioning in secondary lymphoid organs. EF Tfh cells (PD1^low^) produce high levels of IL-21 and low levels of CXCL13 compared with GC PD1^high^ CXCR5^high^ Tfh cells (57). Therefore, IL-21 availability appears as a key element in the initial activation of B cells (58). Our data demonstrated that IFN-α could bypass the IL-21 dependent restriction of PB differentiation in NARB^+^ cells. Early viral responses are mediated by the EF pathway and rely on type I IFN activation (59). Precise components of the EF B-cell responses are still not fully understood, and recently EF effector B cells with pathogenic activity were described in patients with systemic lupus erythematosus expressing CD11c and T-BET. ANav B cells are the precursors of autoreactive CXCR5^-^ CD11c^+^ cells (DN2) and EF PB (60), and their development may be related to an aberrant IFN-γ and IL-21 environment. Our phenotypic analysis showed that NARB^+^ were a distinct subset from the aNav B cells, as they did not express T-BET and seemed not to be chronically activated (normal expression of CD21 and CD19). Those observations were in favor of a physiological non-pathogenic subset; however, we could not assume any functional relationship between the different EF precursor populations and how they might interact. The control of the EF response and the reactivation of natural innate or chronically activated B cells appear essential in autoimmunity and acute viral infections such as SARS-CoV2 or influenza (61). The convergent and balanced existence of different EF progenitors could participate in the homeostasis of a such response, providing protective rapid immunity but which could lead to an exacerbated inflammatory response (9). In this regard, innate MZB cell (USM) differentiation seems impaired in patients with systemic lupus erythematosus and Sjögren’s syndrome (63).Therefore, elucidation of the cellular and molecular actors controlling the initial B-cell activation toward MZB or FO developmental pathway would be promising for better care of systemic autoimmune diseases and hyper-inflammatory syndromes.

## CONTRIBUTORSHIP

M.B, L.L.P, and S.H designed the study, performed experiments, analyzed data, and wrote the paper. M.B, M.M.C, and P.H performed experiments and analyzed data. A.G, CLD and E.P performed RNA sequencing experiments and analyzed data. D.C, C.J, V.D and J-O.P discussed data and edited the manuscript.

## ACKNOWLEDGMENTS

The authors would like to acknowledge the Cytometry Core Facility Hyperion (Brest, France) for their technical assistance. We thank Nadège Marec and Pierre Pochard for the assistance in cell sorting, and Patrice Hémon for the HYPERION and CYTOF experiments.

## FUNDING INFO

This work has been carried out thanks to the support of the LabEx IGO program (n° ANR-11-LABX-0016-01) funded by the «Investissements d’Avenir» French Government program, managed by the French National Research Agency (ANR)”.

## ETHICAL APPROVAL INFORMATION

The study was performed in accordance with the Declaration of Helsinki and was approved by ethical committees.

## DATA SHARING STATEMENT

The data that support the findings of this study are available from the first and the corresponding authors upon request

## BIBLIOGRAPHY

1. Weisel F, Shlomchik M. Memory B Cells of Mice and Humans. Annual review of immunology. 2017;35:255–84.

2. Baker D, Marta M, Pryce G, Giovannoni G, Schmierer K. Memory B Cells are Major Targets for Effective Immunotherapy in Relapsing Multiple Sclerosis. EBioMedicine. 2017;16:41–50.

3. Rajewsky K. Clonal selection and learning in the antibody system. Nature. 1996;381(6585):751–8.

4. Tangye SG, Tarlinton DM. Memory B cells: effectors of long-lived immune responses. European journal of immunology. 2009;39(8):2065–75.

5. Reynaud CA, Descatoire M, Dogan I, Huetz F, Weller S, Weill JC. IgM memory B cells: a mouse/human paradox. Cellular and molecular life sciences : CMLS. 2012;69(10):1625–34.

6. Lee SK, Rigby RJ, Zotos D, Tsai LM, Kawamoto S, Marshall JL, et al. B cell priming for extrafollicular antibody responses requires Bcl-6 expression by T cells. The Journal of experimental medicine. 2011;208(7):1377–88.

7. Cerutti A, Cols M, Puga I. Marginal zone B cells: virtues of innate-like antibody-producing lymphocytes. Nature reviews Immunology. 2013;13(2):118–32.

8. Paus D, Phan TG, Chan TD, Gardam S, Basten A, Brink R. Antigen recognition strength regulates the choice between extrafollicular plasma cell and germinal center B cell differentiation. The Journal of experimental medicine. 2006;203(4):1081–91.

9. Grasseau A, Boudigou M, Le Pottier L, Chriti N, Cornec D, Pers JO, et al. Innate B Cells: the Archetype of Protective Immune Cells. Clinical reviews in allergy & immunology. 2020;58(1):92–106.

10. Recher M, Berglund LJ, Avery DT, Cowan MJ, Gennery AR, Smart J, et al. IL-21 is the primary common gamma chain-binding cytokine required for human B-cell differentiation in vivo. Blood. 2011;118(26):6824–35.

11. Tangye SG, Ma CS. Regulation of the germinal center and humoral immunity by interleukin-21. The Journal of experimental medicine. 2020;217(1).

12. Avery DT, Deenick EK, Ma CS, Suryani S, Simpson N, Chew GY, et al. B cell-intrinsic signaling through IL-21 receptor and STAT3 is required for establishing long-lived antibody responses in humans. The Journal of experimental medicine. 2010;207(1):155–71.

13. Pinto AK, Ramos HJ, Wu X, Aggarwal S, Shrestha B, Gorman M, et al. Deficient IFN signaling by myeloid cells leads to MAVS-dependent virus-induced sepsis. PLoS pathogens. 2014;10(4):e1004086.

14. Durbin JE, Fernandez-Sesma A, Lee CK, Rao TD, Frey AB, Moran TM, et al. Type I IFN modulates innate and specific antiviral immunity. Journal of immunology. 2000;164(8):4220–8.

15. Cervantes-Barragan L, Zust R, Weber F, Spiegel M, Lang KS, Akira S, et al. Control of coronavirus infection through plasmacytoid dendritic-cell-derived type I interferon. Blood. 2007;109(3):1131–7.

16. Herber D, Brown TP, Liang S, Young DA, Collins M, Dunussi-Joannopoulos K. IL-21 has a pathogenic role in a lupus-prone mouse model and its blockade with IL-21R.Fc reduces disease progression. Journal of immunology. 2007;178(6):3822–30.

17. Domeier PP, Chodisetti SB, Schell SL, Kawasawa YI, Fasnacht MJ, Soni C, et al. B-Cell-Intrinsic Type 1 Interferon Signaling Is Crucial for Loss of Tolerance and the Development of Autoreactive B Cells. Cell reports. 2018;24(2):406–18.

18. Zumaquero E, Stone SL, Scharer CD, Jenks SA, Nellore A, Mousseau B, et al. IFNgamma induces epigenetic programming of human T-bet(hi) B cells and promotes TLR7/8 and IL-21 induced differentiation. eLife. 2019;8.

19. Deenick EK, Avery DT, Chan A, Berglund LJ, Ives ML, Moens L, et al. Naive and memory human B cells have distinct requirements for STAT3 activation to differentiate into antibody-secreting plasma cells. The Journal of experimental medicine. 2013;210(12):2739–53.

20. Lam JH, Baumgarth N. The Multifaceted B Cell Response to Influenza Virus. Journal of immunology. 2019;202(2):351–9.

21. Jenks SA, Cashman KS, Zumaquero E, Marigorta UM, Patel AV, Wang X, et al. Distinct Effector B Cells Induced by Unregulated Toll-like Receptor 7 Contribute to Pathogenic Responses in Systemic Lupus Erythematosus. Immunity. 2018;49(4):725–39 e6.

22. Sanz I, Wei C, Jenks SA, Cashman KS, Tipton C, Woodruff MC, et al. Challenges and Opportunities for Consistent Classification of Human B Cell and Plasma Cell Populations. Frontiers in immunology. 2019;10:2458.

23. Hipp N, Symington H, Pastoret C, Caron G, Monvoisin C, Tarte K, et al. IL-2 imprints human naive B cell fate towards plasma cell through ERK/ELK1-mediated BACH2 repression. Nature communications. 2017;8(1):1443.

24. Avery DT, Bryant VL, Ma CS, de Waal Malefyt R, Tangye SG. IL-21-induced isotype switching to IgG and IgA by human naive B cells is differentially regulated by IL-4. Journal of immunology. 2008;181(3):1767–79.

25. Auladell M, Nguyen TH, Garcillan B, Mackay F, Kedzierska K, Fox A. Distinguishing naive- from memory-derived human B cells during acute responses. Clinical & translational immunology. 2019;8(11):e01090.

26. Jourdan M, Caraux A, De Vos J, Fiol G, Larroque M, Cognot C, et al. An in vitro model of differentiation of memory B cells into plasmablasts and plasma cells including detailed phenotypic and molecular characterization. Blood. 2009;114(25):5173–81.

27. Carotta S, Willis SN, Hasbold J, Inouye M, Pang SH, Emslie D, et al. The transcription factors IRF8 and PU.1 negatively regulate plasma cell differentiation. The Journal of experimental medicine. 2014;211(11):2169–81.

28. Pape KA, Taylor JJ, Maul RW, Gearhart PJ, Jenkins MK. Different B cell populations mediate early and late memory during an endogenous immune response. Science. 2011;331(6021):1203–7.

29. Parrish-Novak J, Dillon SR, Nelson A, Hammond A, Sprecher C, Gross JA, et al. Interleukin 21 and its receptor are involved in NK cell expansion and regulation of lymphocyte function. Nature. 2000;408(6808):57–63.

30. Wan CK, Oh J, Li P, West EE, Wong EA, Andraski AB, et al. The cytokines IL-21 and GM-CSF have opposing regulatory roles in the apoptosis of conventional dendritic cells. Immunity. 2013;38(3):514–27.

31. Jin H, Carrio R, Yu A, Malek TR. Distinct activation signals determine whether IL-21 induces B cell costimulation, growth arrest, or Bim-dependent apoptosis. Journal of immunology. 2004;173(1):657–65.

32. Mehta DS, Wurster AL, Whitters MJ, Young DA, Collins M, Grusby MJ. IL-21 induces the apoptosis of resting and activated primary B cells. Journal of immunology. 2003;170(8):4111–8.

33. Ettinger R, Sims GP, Fairhurst AM, Robbins R, da Silva YS, Spolski R, et al. IL-21 induces differentiation of human naive and memory B cells into antibody-secreting plasma cells. Journal of immunology. 2005;175(12):7867–79.

34. Inoue T, Moran I, Shinnakasu R, Phan TG, Kurosaki T. Generation of memory B cells and their reactivation. Immunological reviews. 2018;283(1):138–49.

35. Akkaya M, Kwak K, Pierce SK. B cell memory: building two walls of protection against pathogens. Nature reviews Immunology. 2020;20(4):229–38.

36. Wang S, Wang J, Kumar V, Karnell JL, Naiman B, Gross PS, et al. IL-21 drives expansion and plasma cell differentiation of autoreactive CD11c(hi)T-bet(+) B cells in SLE. Nature communications. 2018;9(1):1758.

37. Kim SY, Volsky DJ. PAGE: parametric analysis of gene set enrichment. BMC bioinformatics. 2005;6:144.

38. Zotos D, Coquet JM, Zhang Y, Light A, D’Costa K, Kallies A, et al. IL-21 regulates germinal center B cell differentiation and proliferation through a B cell-intrinsic mechanism. The Journal of experimental medicine. 2010;207(2):365–78.

39. Linterman MA, Beaton L, Yu D, Ramiscal RR, Srivastava M, Hogan JJ, et al. IL-21 acts directly on B cells to regulate Bcl-6 expression and germinal center responses. The Journal of experimental medicine. 2010;207(2):353–63.

40. Jellusova J, Cato MH, Apgar JR, Ramezani-Rad P, Leung CR, Chen C, et al. Gsk3 is a metabolic checkpoint regulator in B cells. Nature immunology. 2017;18(3):303–12.

41. Piovesan D, Tempany J, Di Pietro A, Baas I, Yiannis C, O’Donnell K, et al. c-Myb Regulates the T-Bet-Dependent Differentiation Program in B Cells to Coordinate Antibody Responses. Cell reports. 2017;19(3):461–70.

42. Pal R, Janz M, Galson DL, Gries M, Li S, Johrens K, et al. C/EBPbeta regulates transcription factors critical for proliferation and survival of multiple myeloma cells. Blood. 2009;114(18):3890–8.

43. Glass DR, Tsai AG, Oliveria JP, Hartmann FJ, Kimmey SC, Calderon AA, et al. An Integrated Multi-omic Single-Cell Atlas of Human B Cell Identity. Immunity. 2020;53(1):217–32 e5.

44. Weisel NM, Weisel FJ, Farber DL, Borghesi L, Shen Y, Ma W, et al. Comprehensive analyses of B cell compartments across the human body reveal novel subsets and a gut resident memory phenotype. Blood. 2020.

45. Koethe S, Zander L, Koster S, Annan A, Ebenfelt A, Spencer J, et al. Pivotal advance: CD45RB glycosylation is specifically regulated during human peripheral B cell differentiation. Journal of leukocyte biology. 2011;90(1):5–19.

46. Descatoire M, Weller S, Irtan S, Sarnacki S, Feuillard J, Storck S, et al. Identification of a human splenic marginal zone B cell precursor with NOTCH2-dependent differentiation properties. The Journal of experimental medicine. 2014;211(5):987–1000.

47. Van Gassen S, Callebaut B, Van Helden MJ, Lambrecht BN, Demeester P, Dhaene T, et al. FlowSOM: Using self-organizing maps for visualization and interpretation of cytometry data. Cytometry Part A : the journal of the International Society for Analytical Cytology. 2015;87(7):636–45.

48. Tipton CM, Hom JR, Fucile CF, Rosenberg AF, Sanz I. Understanding B-cell activation and autoantibody repertoire selection in systemic lupus erythematosus: A B-cell immunomics approach. Immunological reviews. 2018;284(1):120–31.

49. Zeng R, Spolski R, Casas E, Zhu W, Levy DE, Leonard WJ. The molecular basis of IL-21-mediated proliferation. Blood. 2007;109(10):4135–42.

50. Srinivasan L, Sasaki Y, Calado DP, Zhang B, Paik JH, DePinho RA, et al. PI3 kinase signals BCR-dependent mature B cell survival. Cell. 2009;139(3):573–86.

51. Hikida M, Casola S, Takahashi N, Kaji T, Takemori T, Rajewsky K, et al. PLC-gamma2 is essential for formation and maintenance of memory B cells. The Journal of experimental medicine. 2009;206(3):681–9.

52. Viant C, Weymar GHJ, Escolano A, Chen S, Hartweger H, Cipolla M, et al. Antibody Affinity Shapes the Choice between Memory and Germinal Center B Cell Fates. Cell. 2020.

53. Bernasconi NL, Traggiai E, Lanzavecchia A. Maintenance of serological memory by polyclonal activation of human memory B cells. Science. 2002;298(5601):2199–202.

54. Kato Y, Abbott RK, Freeman BL, Haupt S, Groschel B, Silva M, et al. Multifaceted Effects of Antigen Valency on B Cell Response Composition and Differentiation In Vivo. Immunity. 2020;53(3):548–63 e8.

55. Zhao Y, Uduman M, Siu JHY, Tull TJ, Sanderson JD, Wu YB, et al. Spatiotemporal segregation of human marginal zone and memory B cell populations in lymphoid tissue. Nature communications. 2018;9(1):3857.

56. Grimsholm O, Piano Mortari E, Davydov AN, Shugay M, Obraztsova AS, Bocci C, et al. The Interplay between CD27(dull) and CD27(bright) B Cells Ensures the Flexibility, Stability, and Resilience of Human B Cell Memory. Cell reports. 2020;30(9):2963–77 e6.

57. Durand M, Walter T, Pirnay T, Naessens T, Gueguen P, Goudot C, et al. Human lymphoid organ cDC2 and macrophages play complementary roles in T follicular helper responses. The Journal of experimental medicine. 2019;216(7):1561–81.

58. Shlomchik MJ, Weisel F. B cell primary immune responses. Immunological reviews. 2019;288(1):5–9.

59. Lam JH, Smith FL, Baumgarth N. B Cell Activation and Response Regulation During Viral Infections. Viral immunology. 2020;33(4):294–306.

60. Tipton CM, Fucile CF, Darce J, Chida A, Ichikawa T, Gregoretti I, et al. Diversity, cellular origin and autoreactivity of antibody-secreting cell population expansions in acute systemic lupus erythematosus. Nature immunology. 2015;16(7):755–65.

61. Woodruff MC, Ramonell RP, Nguyen DC, Cashman KS, Saini AS, Haddad NS, et al. Extrafollicular B cell responses correlate with neutralizing antibodies and morbidity in COVID-19. Nature immunology. 2020;21(12):1506–16.

62. Tull TJ, Pitcher MJ, Guesdon W, Siu JHY, Lebrero-Fernández C, Zhao Y, et al. Human marginal zone B cell development from early T2 progenitors. The Journal of experimental medicine. 2021;218(4):e20202001.

63. Roberts ME, Kaminski D, Jenks SA, Maguire C, Ching K, Burbelo PD, et al. Primary Sjögren’s syndrome is characterized by distinct phenotypic and transcriptional profiles of IgD+ unswitched memory B cells. Arthritis Rheumatology. 2014;66(9):2558–69.

64. Berg S, Kutra D, Kroeger T, Straehle CN, Kausler BX, Haubold C, et al. ilastik: interactive machine learning for (bio)image analysis. Nature Methods. 2019;16(12):1226–1232.

65. McQuin C, Goodman A, Chernyshev V, Kamentsky L, Cimini BA, Karhohs KW, et al. CellProfiler 3.0: Next-generation image processing for biology. PLoS Biology. 2018;16(7):e2005970.

66. Schapiro D, Jackson HW, Raghuraman S, Fischer JR, Zanotelli VRT, Schulz D, et al. histoCAT: analysis of cell phenotypes and interactions in multiplex image cytometry data. Nat Methods. 2017;14(9):873–876.

