## Supplementary Figure Legends for "IL-21 and IFN-alpha have both opposite and redundant role on human innate precursors and memory B-cell differentiation"

**Fig.S1 Sorted B-cell functional responses in presence of IFN-α or IL-21**

**(A)** Gating strategy and cell sorting verification of the four B-cell subsets (NA, USM, SM and DN). **(B)** Assessment of Ig secretory function by ELISPOT in primed B-cell subsets cultured in the presence or not of IFNα or IL-21. Percentages of IgM-(blue), IgA-(yellow), IgG-(red) secreting B cells from NA, USM and SM-primed B cells. **(C)** Evolution of the percentage of IRF4$^{+}$ IRF8$^{low}$ B cells in primed NA (blue) and primed SM (red) B cells cultured in the presence (dashed lines) or not (solid lines) of IL-21 **(D)** Kinetics of the apoptosis (AnnexinV-FITC positive B cells) in primed NA B cells cultured in the presence (orange line) or not (grey line) of IL-21. **(E)** Mean ± SEM BLIMP1 geometric mean intensity (gMFI) on NA (left) and SM B cells (right) at day 0 (D0) (white) and cultured (primed, dark grey) in the presence or not of IFN-α (blue) or IL-21 (orange). N=7 to 8 (for NA), and N=8 to 10 (for SM) independent experiments, and statistical significance was determined by ANOVA test with Tukey'correction. NA: Naive B cells, USM: Unswitched memory B cells, SM: Switched memory B cells, DN: Double-negative B cells.

**Fig.S2 Transcriptional characterization by RNA sequencing of primed naive B cells in the presence of IL-21**

**(A)** Principal component analysis (PCA), PC1 (x-axis, 54 %) and PC2 (y-axis, 15 %) of gene counts from NA and SM B cells at day 0 (D0) and cultured in the presence or not of IFN-α or IL-21. **(B)** Clustering of up-regulated GO pathways from genes in primed NA in the presence of IL-21 *versus* primed NA only. **(C)** Heatmap of genes from IFN signaling differentially expressed between primed NA B cells in the presence of IFN-α ($NA_{IFN}$) *versus* in the presence of IL-21 ($NA_{IL21}$). **(D)** Venn diagrams of up-regulated genes (left) and down-regulated genes (right) between primed NA signature (primed NA *versus* NA at resting state (day 0), green) and in primed NA in the presence of IL-21 signature (+IL-21) (primed NA + IL-21 *versus* NA at resting state (day 0), yellow), characterizing the IL-21 signature. **(E)** KEGG pathway enrichment in primed NA in the presence of IL-21. **(F)** Volcano plot of DEGs between primed NA B cells in the presence of IL-21 ($NA_{IL21}$) and primed SM in the presence of IL-21 ($SM_{IL21}$). **(G)** Heatmap of important genes differentially expressed between primed NA B cells in the presence of IL-21 ($NA_{IL21}$) *versus* primed SM B cells in the presence of IL-21 ($SM_{IL21}$). NA: Naive B cells, SM: Switched memory B cells.

**Fig.S3 Gating strategy for the identification of B cells from whole blood of healthy donors by CyTOF analysis.**

**Fig.S4 Visualization of $IgD^+$ $CD27^-$ $CD45RB^+$ ($NARB^+$) cells and their neighborhood in tissues by imaging mass cytometry**

**(A)** Expression of Ki67, CD45RB, CD27, IgD and PanKeratin in gut (left), in tonsil (middle) and spleen (right) with Hyperion mass imager. **(B)** Heatmaps of expression density of CCR6, CD138, CD20, CD45RB, IgD, Ki67, CD27, CD4, CD138, FOXP3, PD1, CD8a, CD68, ColI and MPO, PanKeratin and CD8a markers in the 10 meta-clusters obtained with FlowSOM clustering in gut (left), in spleen (middle), in tonsil (right). **(C)** Manual gating of the major cell subsets loaded on the t-SNE plots in gut (left), in spleen (middle), in tonsil (right): polymorphonuclear neutrophils (PMNs), germinal center B cells (GC), $IgD^{+}$ $CD27^{-}$ naive $CD45RB^{+}$ B cells ($NARB^{+}$) **(D)** Representative images of $Ki67^{+}$ B cells, $NARB^{+}$ B cells and CD4 T cells in the tonsil (left), and $CD20^{+}$, $NARB^{+}$, and $CD4^{+}$ in the gut with the heatmap of $NARB^{+}$. neighborood analysis in the gut. dashed line squares identified cell interactions.
