## Supplementary figures and images for "IL-21 and IFN-alpha have both opposite and redundant role on human innate precursors and memory B-cell differentiation"

### Supplementary Figure 1

#### Supplementary Figure 1

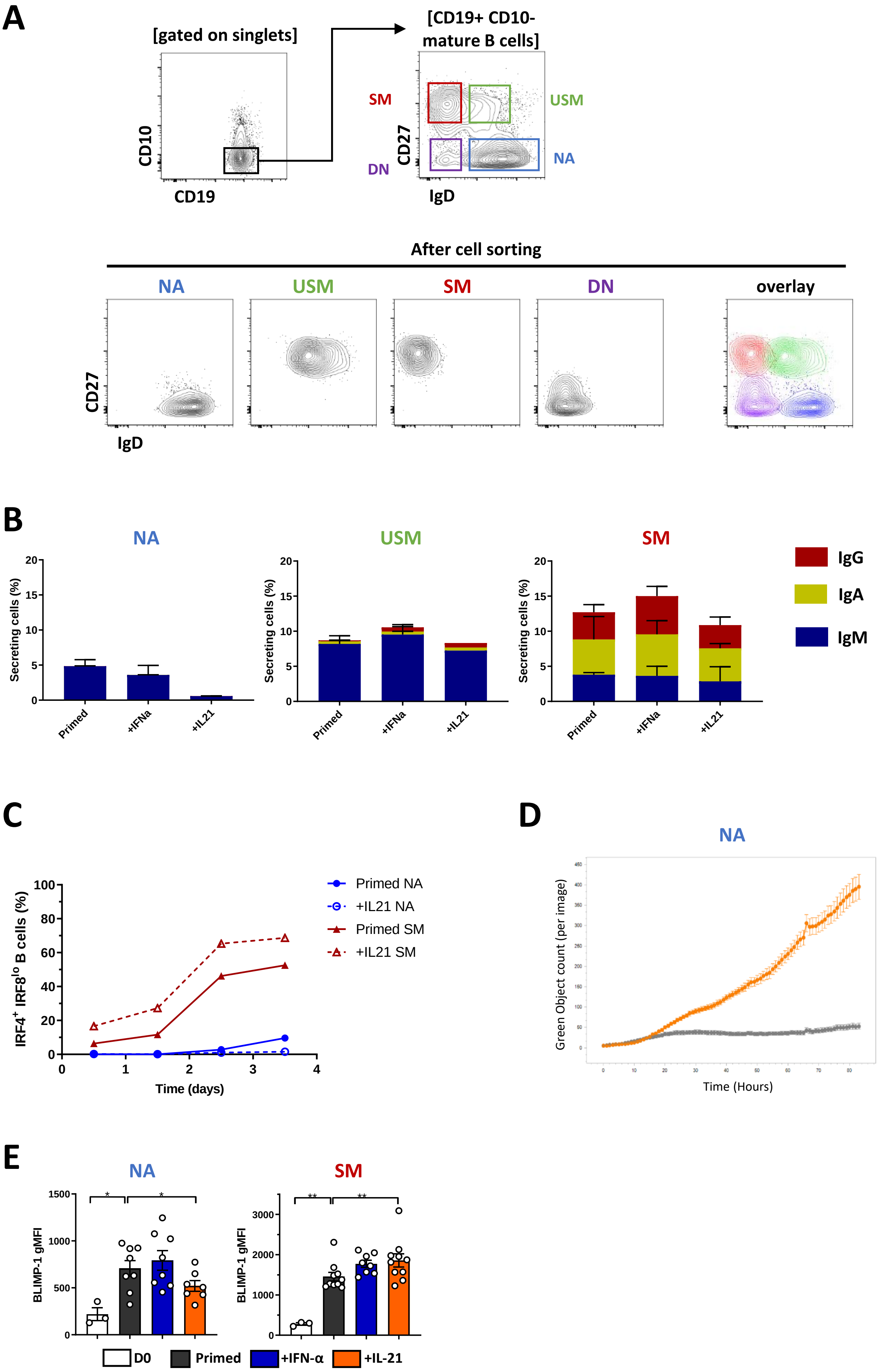

### Supplementary Figure 2

#### Supplementary Figure 2

A

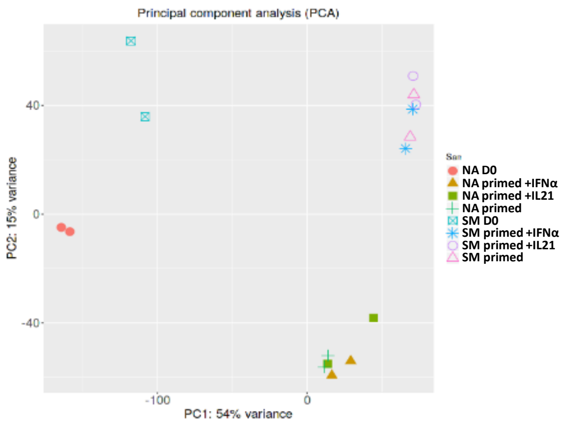


B

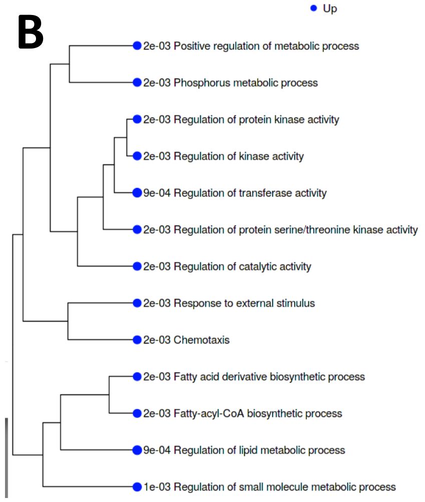


C

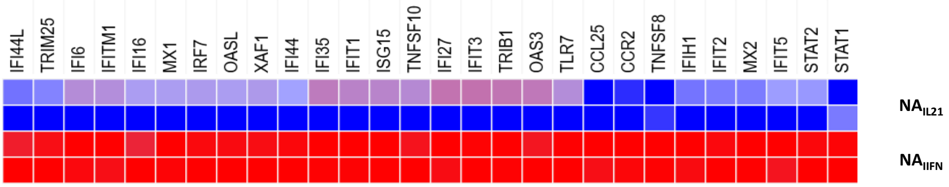


D

| 1,5 fold, 5 %FDR | NA-primed | +IL-21 |
|---|---|---|
| Up-regulated | 4112 | 4563 |
| Down-regulated | 3479 | 3795 |

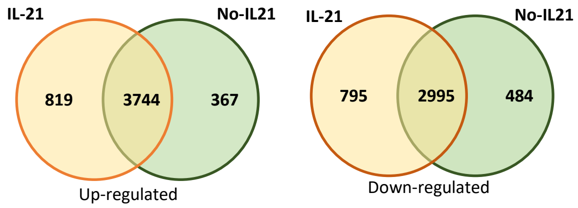


E

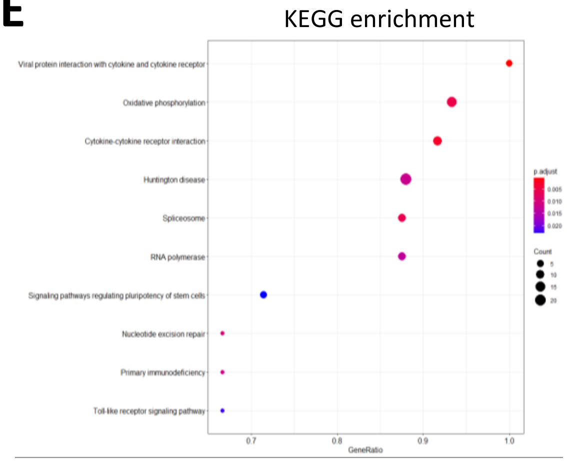


F

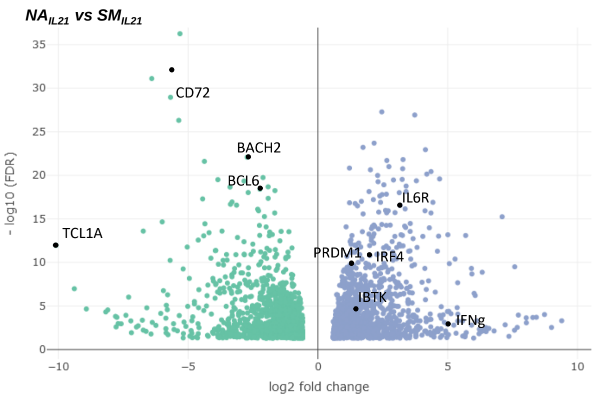


G

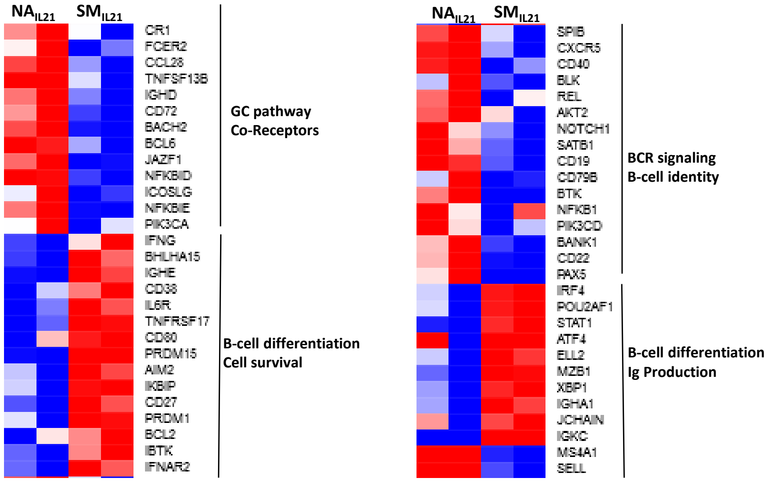

### Supplementary Figure 3

#### Supplementary Figure 3

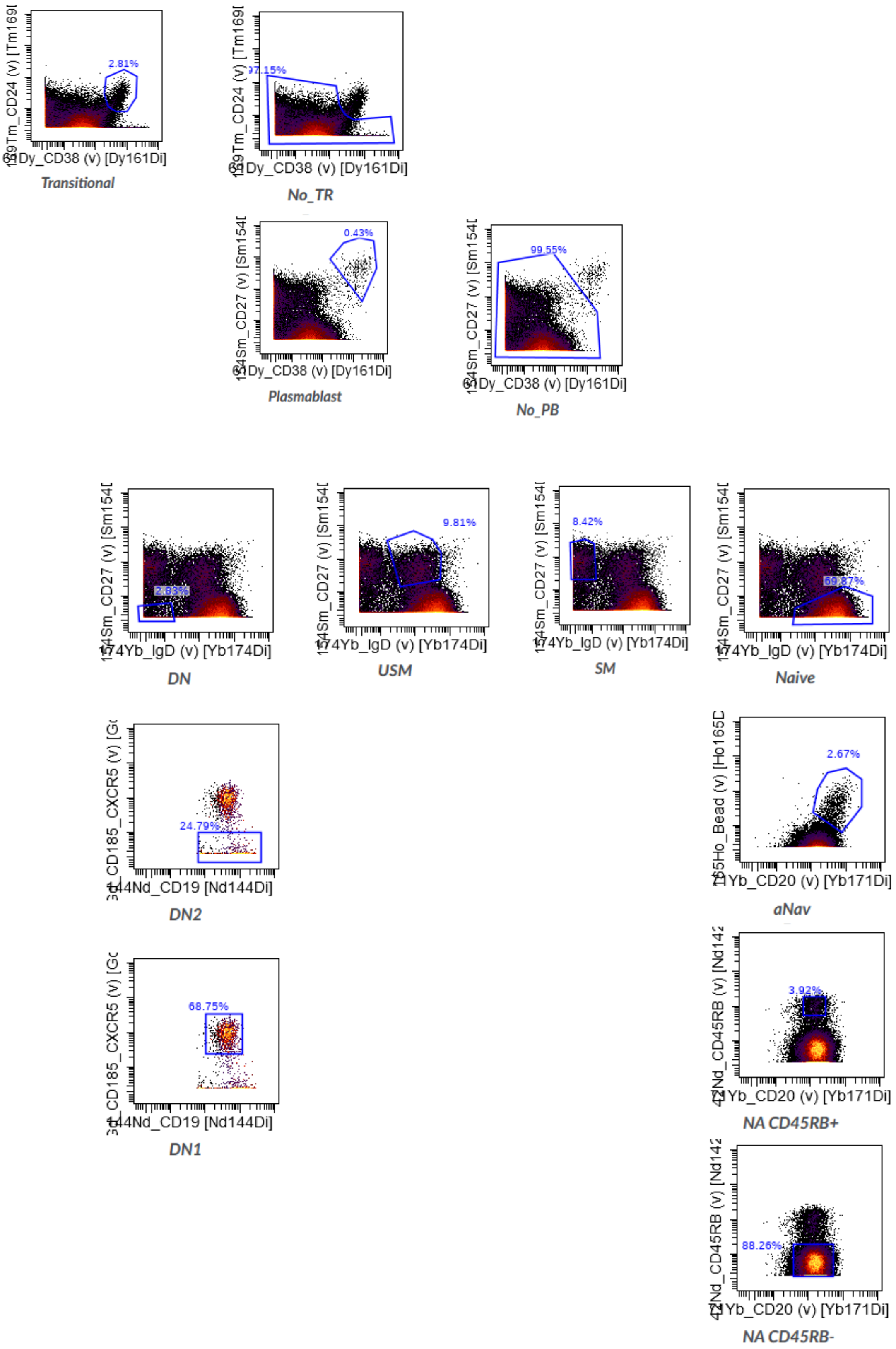

### Supplementary Figure 4

#### Supplementary Figure 4

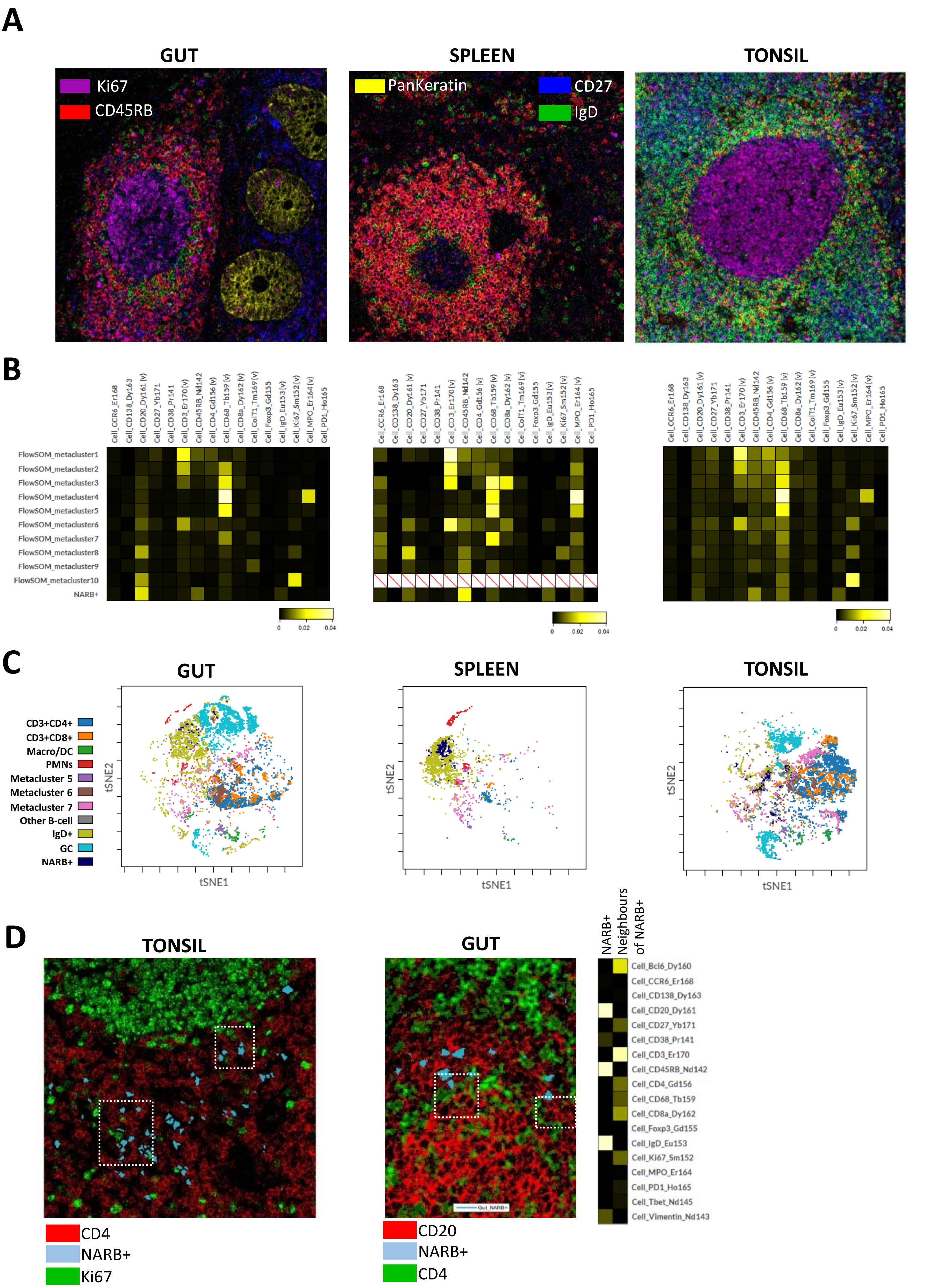
